# Joint profiling of cell morphology and gene expression during *in vitro* neurodevelopment

**DOI:** 10.1101/2023.12.03.569583

**Authors:** Adithi Sundaresh, Dimitri Meistermann, Riina Lampela, Zhiyu Yang, Rosa Woldegebriel, Andrea Ganna, Pau Puigdevall, Helena Kilpinen

**Affiliations:** Helsinki Institute of Life Science (HiLIFE), University of Helsinki, Finland; Institute for Molecular Medicine Finland (FIMM), University of Helsinki, Finland; Faculty of Medicine, University of Helsinki, Finland; Faculty of Biological and Environmental Sciences, University of Helsinki, Finland

**Author notes:** Equal contribution (Adithi Sundaresh, Dimitri Meistermann).

**Keywords:** stem cells, iPSC, differentiation, neurodevelopment, cortical neurons, single-cell genomics, transcriptomics, cell painting, high-content imaging, disease modeling

## Abstract

Differentiation of induced pluripotent stem cells (iPSC) towards neuronal lineages has enabled diverse cellular models of human neurodevelopment and related disorders. However, *in vitro* differentiation is a variable process that frequently leads to heterogeneous cell populations that may confound disease-relevant phenotypes. Here, we report a thorough characterization of cortical neuron differentiation from iPSCs using joint profiling of neuronal morphology and transcriptomics in single cells. We assayed 60,000 developing neurons across three timepoints with CellPainting, a high-content imaging method, and single-cell RNA-sequencing (scRNA-seq). By modeling the relationship between morphological features and gene expression within our differentiation system, we annotated image-based features with biological functions and found that while phenotypes related to cell morphology capture broader neuronal classes than scRNA-seq, they enhance our ability to quantify the biological processes that drive neuronal differentiation over time, such as mitochondrial function and cell cycle. Further, we found that while over 60% of the cells match those seen in the fetal brain, 28% represented metabolically abnormal cell states and broader neuronal classes specific to *in vitro* cells. Finally, we show that specific subtypes of iPSC-derived cortical neurons are a relevant model for a range of brain-related complex traits, including schizophrenia and bipolar disorder, highlighting the potential of multi-modal single cell phenotyping for disease modeling.

## Introduction

Induced pluripotent stem cells (iPSCs) have enabled cell-level profiling experiments in many previously inaccessible cell types and lineages. For example, human iPSC-derived neurons and glia have revolutionized the study of brain-related disorders, which previously relied heavily on animal models and post-mortem tissue. However, iPSC-based differentiation systems are inherently variable^1–3^ and for any given protocol, the full spectrum of cell types generated *in vitro* is not known. For example, *in vitro* conditions can give rise to cell types and states that are not seen *in vivo*^4^. Single-cell RNA-sequencing (scRNA-seq) has transformed the resolution at which iPSC-derived cell types can be characterized. However, gene expression levels alone do not comprehensively reflect changes in cellular function and the biological processes that ultimately drive disease pathophysiology^5^, highlighting the need to measure additional cellular phenotypes.

In this study, we explored the utility of Cell Painting (CP), a high-content image-based assay, in expanding the spectrum of phenotypic readouts available from single cells. We performed joint profiling of cellular morphology (Cell Painting) and gene expression (scRNA-seq) in iPSC-derived cortical neurons. Cell Painting uses fluorescent dyes to label different basic organelles of the cell, such as the nucleus, mitochondria, and the endoplasmic reticulum (ER) from which hundreds of image-based features can be derived, representing the morphological profile of each cell^6,7^. Compared with other imaging technologies, CP is more affordable and transferable across cell types. It has been used extensively for large-scale screening experiments, for e.g. in the context of drugs^8–10^, and therefore is highly suitable for the unbiased characterization of *in vitro* systems where phenotypes are not known *a priori*.

We applied Cell Painting to an established cortical neuron differentiation system based on dual- SMAD inhibition, chosen for its reported ability to recapitulate the progression of neurodevelopment *in vitro*^11,12^. We collected data from >60,000 cells from four healthy donors across three time points corresponding to early progenitors (day 20), intermediate progenitors (day 40) and maturing cortical neurons (day 70) (**Fig. 1a**). Cell type heterogeneity in this system, and in the brain more generally, is high, and our aim was to test to what degree CP is able to disentangle closely related cell types and what phenotypic information is gained beyond transcriptomic data. To establish a cell type reference, we also collected transcriptomic data from the same developmental time points across the differentiation and compared them to fetal cell types *in vivo* (**Fig. 2a**). CP data is typically analyzed at well-level and little has been done to interpret image-derived features directly or in the context of other data modalities to date^13^. Here, we analyzed the CP data at a single-cell resolution and, by leveraging a predictive model^9^, provide links between image features and gene expression levels in developing cortical neurons (**Fig. 2b**). Finally, we evaluated the disease-relevance of the generated cell types using stratified LD Score Regression^14^.

**Figure 1:**
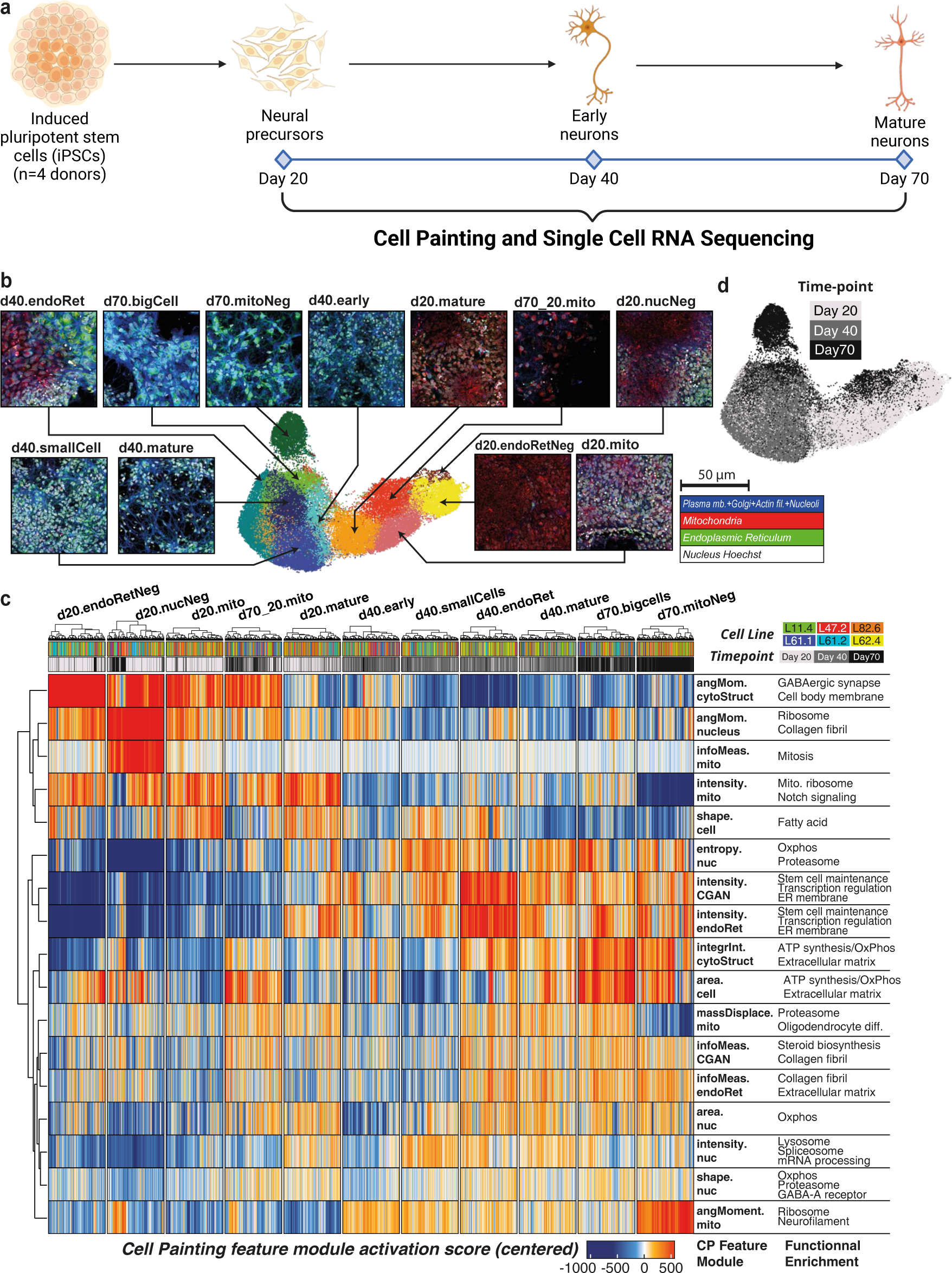
Overview of the experimental design and Cell Painting at the single-cell level. **(a)** Overview of the experiment. 6 iPSC om 4 donors were differentiated to cortical neural fate and assayed at days 20, 40 and 70 of the differentiation using Cell Painting RNA-seq. **(b)** UMAP of the Cell Painting dataset at the single-cell level. Cells are colored and labeled by unsupervised Leiden ng. A composite image representative of cells belonging to each CP cluster is indicated by arrows (staining legend bottom-right). tmap of CP feature-module activation scores. 541 cells (minimum cluster size) were drawn from each CP cluster to ensure an equal ution of each CP cluster to the heatmap. The CP feature modules represent sets of CP features that are highly correlated. Right- olumn contains selected highly-enriched terms from GSEA of CP feature modules. **(d)** Differentiation day (timepoints) projected on UMAP.

**Figure 2:**
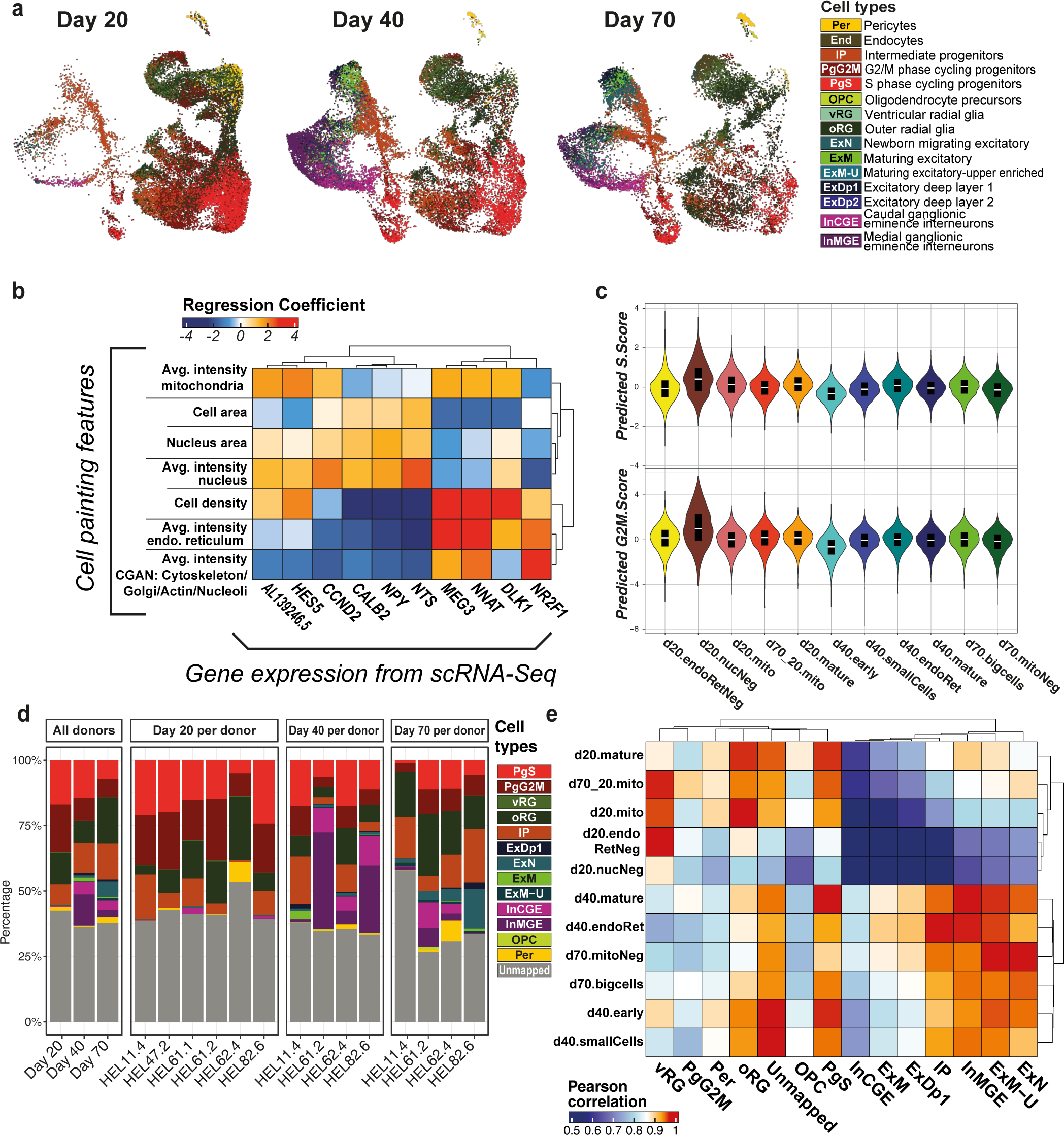
Joint profiling of neural cell types observed *in vitro*. **(a)** UMAP of cell types as identified by scRNA-seq analysis, split by int. **(b)** Representative heatmap of the model linking expression of marker genes (rows) to CP image-based features (columns). lin plots of predicted cell cycle scores per cell painting cluster. **(d)** Cell type composition from the scRNA-seq dataset. The n the left represents aggregated cell type proportions across donors per time point, while the other facets represent individual nts, depicting cell type proportions for each donor. **(e)** Correlation heatmap of cell types to cell painting clusters, between average eatures values per cell clusters and average of predicted CP features from scRNA-seq. **Abbreviations:** PgS - S phase progenitors, - G2M phase progenitors, vRG - ventral radial glia, oRG - outer radial glia, IP - intermediate progenitors, ExDp1/2 - Excitatory deep eurons 1/2, ExN - mirating excitatory neurons, ExM - maturing excitatory neurons, ExM-U - upper-layer enriched maturing excitatory , InCGE - caudal ganglionic eminence interneurons, InMGE - medial ganglionic eminence interneurons, OPC - oligodendrocyte ors, Per - pericytes, End - endothelial cells.

We found that over 60% of the cells in our cortical system reliably matched cell types in the fetal brain, while an additional 28% represented broader neuronal classes observed in stem cell- derived organoids and metabolically abnormal cell states. While CP alone was able to resolve only the larger supertypes of cells (progenitors versus neurons), the joint analysis of cell morphology features and gene expression recapitulated known cell biology, as previously reported^15^. Donor-specific changes were captured by both assays, and image-based features were informative in quantifying the dynamic nature of cellular differentiation, as observed with changes in mitochondrial intensity and cell cycle over time. However, our results suggest that to characterize a heterogeneous system, such as neurons, based on cell morphology requires targeted development of the CP assay for the cell types of interest^16^. Overall, we found that specific subtypes of iPSC-derived cortical neurons capture a significant fraction of heritability of schizophrenia and bipolar disorder and multimodal phenotyping of the most relevant cell populations holds much promise for identifying actionable biological processes. To this end, we provide here a single cell atlas of normal 2D cortical differentiation and a proof-of-concept approach to integrate transcriptomic and image-based phenotypes from single cells for subsequent disease modeling studies.

## Results

### 1. Cell Painting at the single-cell level captures developmental progression

Cellular organelles such as mitochondria and the ER are known to play an important role in neurodevelopment, specifically in meeting dynamic energetic and metabolic requirements^17^. Aiming to characterize the morphological features of developing neurons *in vitro*, we differentiated iPSCs from four donors towards cortical neuronal fate and characterized them at days 20, 40 and 70 of the differentiation (**Fig. 1a, Tables 1, S1-S4**). As a preliminary benchmarking of whether the three timepoints captured the expected cell types, we confirmed expression of canonical markers of neural progenitors (*NESTIN*), intermediate progenitors (*EOMES*) and neurons (*TUJ1*) via immunocytochemistry^11^ (**Fig. S1a**) (**Tables S5-6, Methods**). To extend these morphological profiles in an unbiased manner, we profiled the same time points using Cell Painting. We obtained CP image-based features such as fluorescence intensities, texture and cell shape measures from each CP channel, which together represent the overall morphological profile of the cell. These features were extracted into a matrix per single cell and, after quality control and regularization (**Methods**), we obtained a feature matrix of 223 features from 54,415 cells. Based on observations from performing well-to-well correlation, we found that differentiation day was, as expected, the major source of variation (**Fig. S2a**).

To maintain and explore heterogeneity in the dataset, the feature matrix was then analyzed in a similar manner to single cell RNA sequencing (scRNA-seq) data. Leiden clusters projected on a Uniform Manifold Approximation and Projection (UMAP) (**Fig. 1b**) were used to explore the main phenotypic states captured by Cell Painting. We found that the unsupervised clusters captured variation in both cell size and organelle staining intensities. In light of the high modularity of CP features, we defined feature modules based on the correlation between CP features (**Figs. 1c**, **S2b**). Correlated features within the same module usually corresponded to multiple aspects of the same measurement, for example cell area and perimeter, or the average and minimum intensity of each channel. Interestingly, there was a global correlation between the channel comprising the cell membrane, Golgi apparatus, actin cytoskeleton and nucleoli (CGAN) and the ER channel. In contrast, some pairs of Cell Painting features were highly anticorrelated (**Fig. S2b**). In our study, this was observed between cell size and density, suggesting that smaller cells are more likely to be found in areas of higher density (**Fig. S2c**). Furthermore, a near-perfect negative correlation was detected between texture values related to the uniformity of the channel (‘Angular Second Moment’) and the intensity within each channel. This phenomenon suggests that these two measurements may represent inverse aspects of the same underlying characteristic.

The distribution of feature values shows that despite the difference between timepoints, CP was able to capture a relative continuity across them. Clusters composed mainly of day 20 cells were characterized by high intensities for the mitochondria channel (**Fig. 1c-d**), and clusters with mainly day 40 cells by medium mitochondria intensity and high ER intensity. Interestingly, day 70 was the most heterogeneous timepoint with three distinct profiles: cells with a large area that were close to d40 cells in terms of features value (‘d70.bigCells’), cells with very low mitochondria intensity (‘d70.mitoNeg’) and cells that clustered with d20 cells (‘d70_20.mito’).

### 2. Biological annotation of image features reveal separation of broad neuronal cell types

To improve the interpretability of CP features and link them to gene expression, we additionally profiled the transcriptomes of developing neurons from the same four donors at days 20, 40 and 70 of the differentiation using scRNA-seq. We then modeled the relationship between Cell Painting and scRNA-seq similarly to Haghighi *et al.*^9^, by using the common experimental design between the two assays to train lasso regression models that can be used to predict the corresponding CP feature values of the scRNA-seq dataset (**Fig. 2b, Methods**). We used the coefficients matrix from the resulting model to perform a functional enrichment of previously identified CP feature modules, using the median of gene coefficients per feature module as an input to Gene Set Enrichment Analysis (GSEA) (**Fig. 1c**, **Supplementary Data 1**)^18,19^. The analysis revealed a significant redundancy among the enriched terms observed, with a predominant focus on the proteasome, extracellular matrix, and mitochondrial respiration. This pattern underscores the property of Cell Painting to predominantly capture variation related to cell morphology rather than specific pathway regulation. In most cases, the enriched terms per module were closely linked to the module-associated features, increasing the confidence in our model. For example, in a module composed of features associated with the ER channel, the most enriched terms were related to the ER membrane. Interestingly, other general terms such as intensity of mitochondria were more linked to mitochondrial ribosomes than mitochondrial respiration or oxidative phosphorylation.

A small pool of d20 cells (‘d20.nucNeg’) was particularly linked with mitosis. To further explore the possible link between CP and cell cycle, we predicted the cell cycle scores (G2M score, S score) in the CP dataset by using the predicted CP features matrix on scRNA-seq profiled cells and their annotated cell cycle scores (**Methods**). Using our regression model, ‘d20.nucNeg’ cells showed the highest median values for both G2M score and S score (**Fig. 2c**), and, coupled with their particular morphology, suggests that CP is able to capture cycling cells at least at day 20.

Neuronal differentiation is a tightly regulated process, generating heterogeneous cell types in a temporal manner. To resolve cell types across the three time points of the Cell Painting dataset, we first classified the cell types of the 62,929 cells present in our scRNA-seq dataset. It has been established previously that *in vitro* generated neurons more closely resemble fetal rather than adult neuronal cell types^20,21^. We thus used the reference mapping approach from the Seurat package^22^ to transfer cell type labels from a well-annotated mid-gestation (gestational weeks 17- 18) fetal reference^23^ onto our query dataset, setting a mapping quality threshold of 0.5 (**Methods, Fig. S3a-b**). With this approach, we annotated 60% of our cells with high confidence and without being limited to a few canonical markers, aiming to better describe the dynamics of *in vitro* cortical differentiation. We identified 14 cell types produced across the three time points, capturing a large portion of the heterogeneity seen in developing neurons (**Fig. 2a,d**). These include various progenitor cell types (cycling progenitors, ventral and outer radial glia) as well as intermediate progenitors and maturing neuronal cells (deep layer and maturing upper layer excitatory neurons). We also detected inhibitory neurons within our cortical system (MGE- and CGE-like interneurons), as has been reported recently^24^.

We used the predicted CP feature matrix to assess the correlation between CP clusters and the cell types inferred from scRNA-seq (**Fig. 2e**). We found that CP captured broader cell supertypes, with each CP cluster correlating to multiple cell types, but distinguishing neuronal cell types from progenitor-like cells. However, the low specificity of this correlation did not enable distinguishing individual cell types, demonstrating the efficacy of CP in classifying these two broad categories of cells, yet highlighting its limitations in differentiating subclasses of neurons.

### 3. Neurodevelopment *in vitro* follows known trajectories and recapitulates cell type heterogeneity seen *in vivo*

Neurodevelopment *in vivo* is characterized by a tightly regulated developmental trajectory, from neural progenitors to radial glia and, via intermediate progenitors, to maturing neurons. Given the limitations in cell type resolution using CP, we sought to instead assess the morphological changes along this trajectory. Using the scRNA-seq data, we first estimated the pseudotime of our differentiating cells across the three time points (**Methods**) and found that the developing cells *in vitro* follow a similar trajectory as seen *in vivo* (**Fig. 3a**). Additionally, by computing gene modules changing as a function of pseudotime with the Monocle3 toolkit, we identified gene sets linked to the development of individual cell types (**Fig. S4a**). These gene sets were enriched for many expected biological processes, such as ‘nuclear division’ in PgG2M cells and ‘ribonucleoprotein complex biogenesis’ in PgS (**Fig. S4b-f**). The IP-associated module was tagged by terms such as ‘cell fate commitment’ and ‘channel activity’, indicative of the transitory role these cells play between progenitors and neurons. Neuronal modules were characterized by formation and regulation of synapses as well as ion channel activity, with the inhibitory neuron-associated module linked to GABAergic signaling.

**Figure 3:**
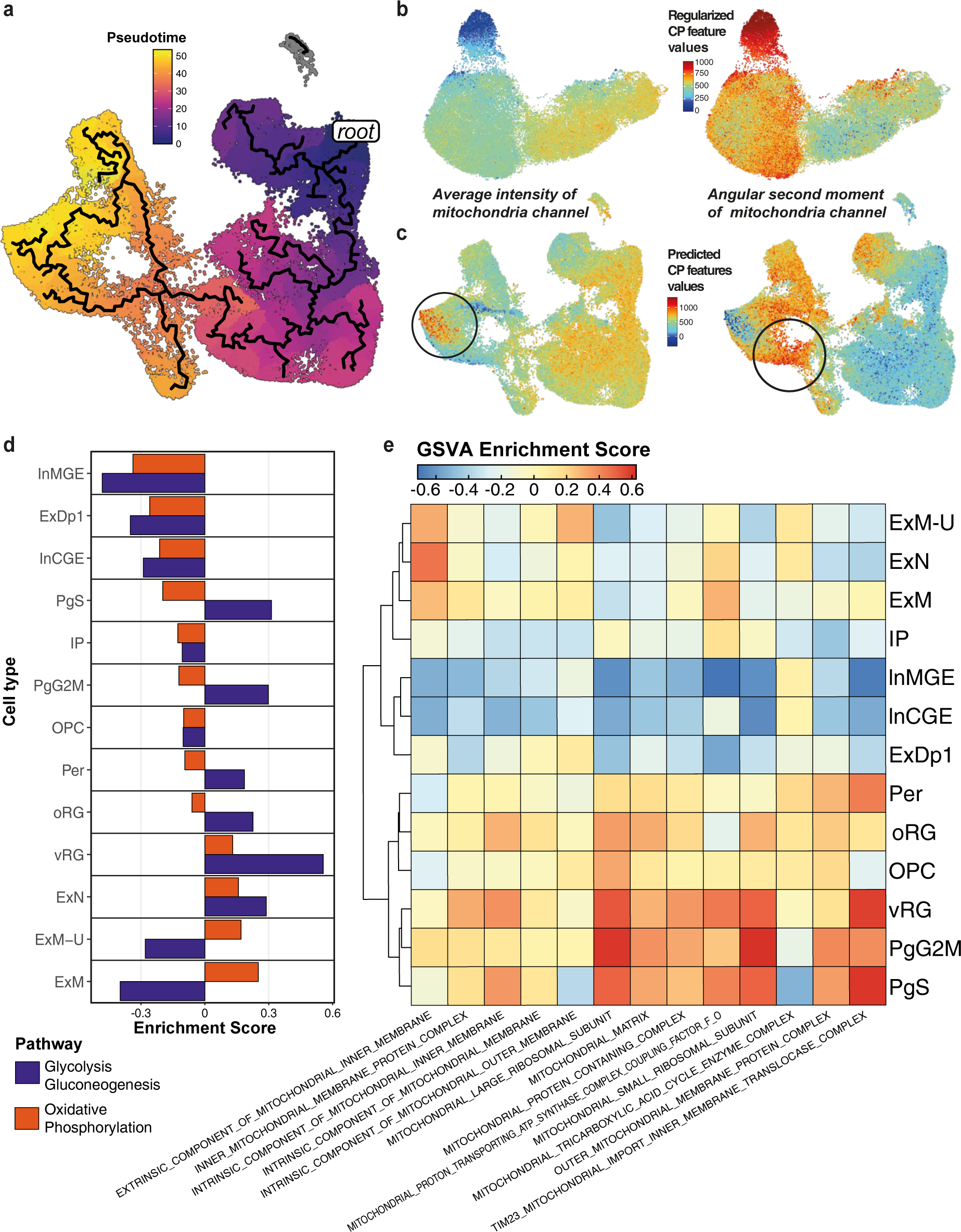
Mitochondrial feature dynamics in Cell Painting and gene expression. **(a)** Predicted pseudotime trajectory across time- n scRNA-seq showing progression from radial glia and progenitors, via intermediate progenitors, towards maturing excitatory and y neurons. Pseudotime is rooted in the predicted earliest node within day 20. **(b)** Feature values for mitochondrial intensity (left) tochondrial texture (right) projected onto the CP UMAP. **(c)** Predicted values for the CP features mitochondrial intensity (left) and (right) projected onto the scRNA-seq UMAP. **(d)** GSVA enrichment scores for glycolysis and oxidative phosphorylation pathways udobulked cell type. Enrichment scores indicate a higher activation score of OxPhos in excitatory neurons, except for the deep subtype (ExDp1). **(e)** GSVA enrichment scores for mitochondria-associated GO terms per pseudobulked cell type.

From the CP data, we observed a distinct pattern of mitochondrial features across the UMAP, with mitochondrial intensity-related features more prominently associated with day 20 cells and decreasing towards day 70, whereas mitochondrial uniformity features (‘angular second moment’) followed the opposing trend (**Fig. 3b**). To associate this trend with the developmental trajectory, we projected these CP features onto the scRNA-seq UMAP confirming that the trend is maintained (**Fig. 3c**); mitochondrial intensity is higher amongst progenitor cell types, whereas uniformity is higher amongst mature/maturing neurons, representing uniform (low intensity) texture. The notable exception is the region of the UMAP occupied by inhibitory neurons, specifically MGE-like interneurons, which instead have high mitochondrial intensity and low uniformity.

Given the cell-type specificity of mitochondrial CP features, we sought to determine whether we could detect changes in the transcription profile linked to mitochondrial metabolism. It has been reported previously that NPCs meet their energetic requirements primarily through glycolytic pathways, and the switch to neuronal fate is associated with a switch to dependency on oxidative phosphorylation (‘OxPhos’)^25^. Gene set variation analysis (GSVA) on pseudo-bulked cell types for both glycolysis and OxPhos (**Methods**) confirmed that most neuronal cell types had lower glycolytic dependency, with all excitatory cell types showing maximum activation for OxPhos except for a subset of excitatory deep layer neurons (‘ExDp1’) (**Fig. 3d**). We further tested for enrichment of gene ontology (GO) terms associated with mitochondria in the gene expression patterns per cell type, finding a clear link between the progenitor cell types and mitochondrial ribosomal subunits, whereas excitatory neuronal cell types were linked to mitochondrial membrane components, known to be linked to OxPhos (**Fig. 3e**). These findings complement the trend observed from the functional enrichment of CP feature modules, where day 20 clusters were enriched in mitochondrial ribosome and mitosis, day 40 in transcriptional regulation, and day 70 in oxidative phosphorylation, overall emphasizing the complementarity of the morphological screen to transcriptomic phenotypes (**Fig. 1c**).

### 4. Donor-specific effects drive neuronal cell fate determination and are captured by both modalities

Inhibitory GABAergic interneurons play a key role in cortical circuits by balancing the excitatory/inhibitory activity and by regulating the formation of synapses. Abnormalities in interneuron development, function or migration have been associated with autism spectrum disorders^26,27^ and other neurodevelopmental disorders (NDDs)^28,29^. Of the neuronal cells identified in our dataset (21% of all cells), nearly 40% were annotated as inhibitory interneurons of either caudal or medial ganglionic eminence (InCGE/InMGE).

Although these have canonically been described to originate from brain regions outside of the cortex, *in vitro* studies have previously reported the generation of GABAergic neurons from cortical progenitors^30,31^. A recent study from Delgado *et al.* confirmed via clonal lineage tracing studies that a subpopulation of cortically born GABAergic neurons was transcriptionally similar to ventrally-derived cortical interneurons, but instead arose from cortical progenitors^24^. The InCGE cells from our dataset were characterized by expression of the *DLX* family of genes, as well as *GAD2, DCX* and *MEIS2*. InMGE cells expressed a subset of canonical markers such as *SST, DCX, MEIS2* but, similar to Delgado *et al.*, differed from true ganglionic eminence interneurons in lack of expression of *LHX6* and *NKX2.1* (**Fig. S5a**). The production of interneurons in our system is valuable to better recapitulate human fetal development considering the relevance of the inhibitory component in neural circuits.

The close clustering of these interneurons with excitatory neuronal cell types prompted us to explore the pathways that differed between them. Previous studies both in mice^32^ and *in vitro* cultures^31^ have indicated the interplay of WNT and SHH signaling as a key factor in maintaining the excitatory/inhibitory balance. However, GSVA analysis of KEGG genes associated with these pathways did not show a consistent upregulation of WNT signaling in all excitatory cell types, nor vice versa with Hedgehog signaling for inhibitory cell types (**Fig. S5b-c**). Instead, we found lower values of enrichment scores for oxidative phosphorylation pathway-associated genes in inhibitory than in excitatory neurons (**Fig. 3d**, also seen from GSEA, **Fig. S5d**).

Additionally, as noted earlier, MGE-like interneurons showed marked differences from the other neuronal cell types in CP-based mitochondrial features (**Fig. 3c**). A single donor (HEL61.2) presented an excess of this inhibitory neuronal subtype across time points (**Fig. 4a**). This same donor was overrepresented in the CP cluster ‘d20.endoRetNeg’, characterized by lowered intensity of the ER channel, and underrepresented in the ER-intense ‘d40.endoRet’ cluster (**Fig. 4b**), pointing towards donor-specific ER effects potentially linked to inhibitory cell fate. Interestingly, we also observed an effect of lowered ribosome-associated genes in gene expression data - the two donors accounting for 80% of inhibitory cell types at days 40 and 70 showed an overall decrease in the KEGG ribosome pathway at these timepoints (**Fig. 4c**). Complementarily, a GSVA of mitochondria-associated GO terms across our *in vitro* cell types (**Fig. 3e**) revealed an overall depletion of mitochondrial ribosome associated genes in inhibitory neurons compared to other cell types, as has been previously reported^33^.

**Figure 4:**
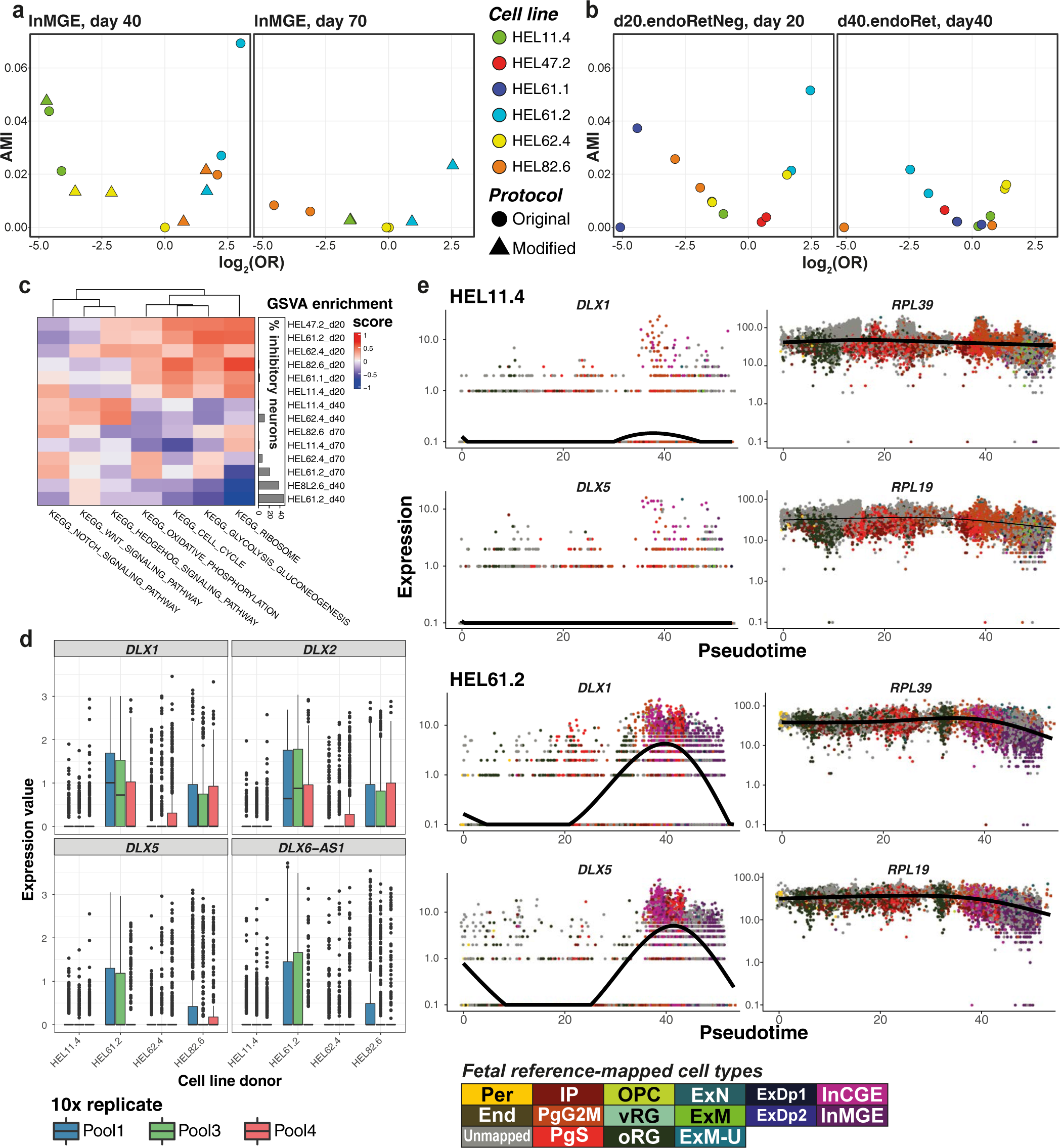
Mechanisms of donor-specific inhibitory neuron production. **(a)** Association plots from scRNA-seq data within InMGE days 40 and 70. Each point represents aggregate expression per technical replicate, donor and the differentiation protocol **b)** Association plots from CP data within the d20.endoRetNeg cluster at d20 and d40.endoRet at day 40. Each point represents ate expression of all wells per induction replicate and donor. **(c)** GSVA enrichment scores from curated KEGG pathways between ental populations (donor x timepoint) with hierarchical clustering on both axes. Annotation barplot to the right of the heatmap the percentage of cells produced by each experimental population annotated as inhibitory cell types (InCGE, InMGE, InhN-O). **(d)** e expression per donor (log-normalized values) of inhibitory interneuron-associated transcription factors within the progenitor pool vRG+PgG2M+PgS) at day 40. The expression per donor is grouped per 10x sample. **(e)** Expression of interneuron-associated enes and ribosomal subunits *RPL39* and *RPL19,* along pseudotime in two donors shown to produce a high (HEL61.2, left) and a L11.4, right) proportion of inhibitory neurons, respectively. Point colors represent fetal-annotated cell types.

In order to elucidate which genes are responsible for the bifurcation between the excitatory and inhibitory fate in our *in vitro* differentiation, we focused on the branch point along the pseudotime trajectory that either produces inhibitory interneurons or IPs, which in turn give rise to excitatory neurons. Within this branch point, we identified modules of genes that were differentially expressed as a function of the pseudotime (**Methods**). As expected, the module marking the inhibitory branch (Module 12, **Fig. S5e**) consisted of TFs known to drive interneuron fate such as the *DLX* family of genes, which was overexpressed in the donor HEL61.2 in day 40 progenitors (**Fig. 4d**). In donors producing inhibitory neurons, expression of the *DLX* family increased with pseudotime, peaking at interneuron production. Concordantly, the expression of genes driving the differential activation of the KEGG ribosome pathway (*RPL39* and *RPL19*) also decreased in donors producing inhibitory neurons (**Fig. 4e**). Altogether, our data is indicative of lowered ribosomal activity and mitochondrial differences between excitatory and inhibitory neurons overall.

### 5. Metabolic differences shape high versus low quality cells produced *in vitro*

It is well established that *in vitro* cell types do not completely resemble their *in vivo* counterparts. When mapped to the reference dataset, 60% of our cells mapped at high confidence to fetal cell types, while the remaining 40% remained ‘unmapped’. These unmapped cells were found to be present across the UMAP (**Fig. 5a**), likely indicating that cells did not fully resemble the fetal transcriptomes and/or were transitioning between cell types, rather than being a single missing or unannotated cell type. By assigning the highest scoring cell type label to each cell in the unmapped fraction of cells, we divided our dataset into high and low quality (HQ/LQ) cells for each annotated cell type (**Methods**). The LQ cells still expressed canonical markers of the cell type they were closest to, although at lower levels, except in the case of outer radial glia (‘oRG’) and intermediate progenitors (‘IP’) (**Fig. S6a**). Additionally, LQ cells did not show consistent differences in pseudotime scores as compared to HQ cells of the same cell type (**Fig. S6b**), which would have been indicative of transitioning cell states.

**Figure 5:**
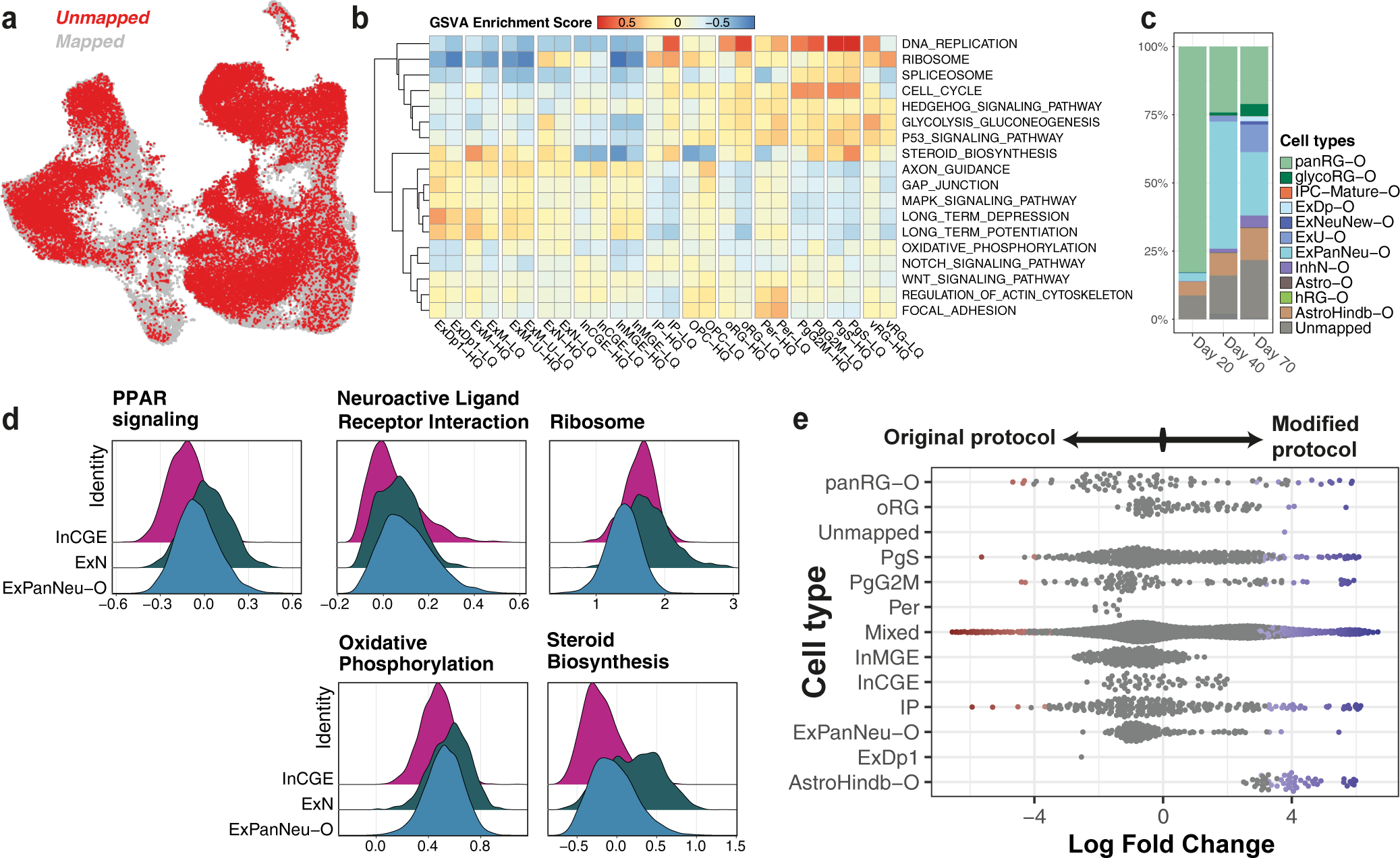
Characterization of low-quality cells within the scRNA-seq dataset. **(a)** UMAP of scRNA-seq data across timepoints mapped cells highlighted in red. **(b)** Enrichment scores from GSVA (across cell types) of key pathways driving the differences n high- and low-quality subsets per pseudobulked cell type. **(c)** Percentage of cells from the initial unmapped fraction that align noid cell types (Bhaduri *et al.*) per time point. **(d)** Ridge plots representing the distribution of module activation scores of key tially activated pathways between pan-neuronal (ExPanNeu-O), excitatory (ExN) and inhibitory (InCGE) neuronal cell types. **(e)** ntial abundance of cell types produced by the original (left) or the modified (right) versions of the differentiation protocol, computed R. **Abbreviations:** panRG - pan radial glia, glycoRG - glycolytic radial glia, IPC-Mature - mature intermediate progenitors, ExDp - ry deep layer, ExNeuNew - newborn excitatory neurons, ExU - excitatory upper layer, ExPanNeu - pan-neuronal (excitatory), InhN tory neurons, Astro - astrocytes, hRG - hindbrain radial glia, AstroHindb - hindbrain astrocytes. “-O” differentiates organoid cell notations from the fetal.

Given this, we assessed what transcriptomic features were driving their lower mapping scores. By comparing differential activation of developmentally relevant pathways between high and low quality cells per cell type, we identified those that differed (**Fig. 5b**). Importantly, although metabolic processes seemed to be implicated overall, we did not see consistent changes across all cell types, rather observing specific changes driving the LQ version of each cell type, such as DNA replication and glycolysis in progenitor cell types (vRG, oRG, IP). Also, we observed the highest steroid biosynthesis activation in mapped maturing excitatory neurons (‘ExM’) compared to unmapped, but the opposite trend was observed within progenitors (PgS, PgG2M).

Aberrant cellular metabolism has previously been reported in other *in vitro* systems, often highlighted in the oxidative phosphorylation and glycolytic pathways^34,35^. In cortical organoids, it has been seen that a fraction of cells does not recapitulate distinct fetal cellular identities^4^.

Hypothesizing that our unmapped cells represented a similar fraction, we annotated our unmapped cells to cortical organoid cell types from *Bhaduri et al.*^4^ using the same label-transfer method and threshold described above (**Methods**). We found that more than 50% of the previously unmapped cells could be attributed to ‘pan-radial glial’ or ‘pan-neuronal’ cell types across all three timepoints (**Fig. 5c**). These were described in the original publication as broad progenitor or neuronal cell classes that did not express features distinctive to any one fetal subtype.

Next, we compared the differentially activated pathways between cells of the closest cell types that passed this bottleneck of subtype acquisition to those that maintained a pan-cell identity. Similar to differences observed between high and low quality progenitor cell types, pan-radial glia differed in ribosome-associated pathways. Pan-neuronal cells, however (‘ExPanNeu-O’), were characterized by features of both excitatory and inhibitory neurons, differing again metabolically such as in oxidative phosphorylation or steroid biosynthesis. We identified modules of the top genes driving differentially activated pathways in these cell types and, by scoring these gene modules per single cell (**Methods**), assessed their distribution in multiple neuronal cell types (**Fig. 5d**). We found that overall, pan-neuronal cells appear to be intermediate to either excitatory or inhibitory neuron specification, except for the ribosome pathway.

Additionally, with the inclusion of the second step of annotation, we identified cell types that were initially missed, mainly astrocytes (**Fig. 5c**). While expected to be produced in this protocol, they were absent in the fetal reference, and thus, the first annotation step alone did not identify them. Of note, our experimental workflow involved the use of two versions of the differentiation protocol (original/modified, see **Methods**), differing in enzyme usage for cell dissociation (see **Table S2** for details). To check for potential effects of the protocol on the generation of cell types, we used *miloR*^36^ to test for differential abundance of annotated cell types between the protocol versions. We found that the astrocyte population (‘AstroHindb-O’) was overrepresented in the modified version of the protocol at day 40 (**Fig. 5e**).

### 6. *In vitro* derived cell types capture heritability of brain-related traits

Finally, to evaluate the relevance of our *in vitro* cortical neurons for disease modeling, we used stratified linkage disequilibrium (LD) score regression^14^ to quantify how much heritability of common diseases and other traits is enriched within genes that are markers of the cell types generated *in vitro*. We analyzed GWAS summary statistics from a set of 79 common traits including 12 brain-related phenotypes (**Table S7, Fig. S7a-b**) (**Methods**). There was a clear enrichment of significant brain-related associations when accounting for all tests (Fisher Test, p=9.617·10^-13^). Comparison of traits captured by *in vitro* cell types (**Fig. 6a**) to those captured by GTEx brain tissues^37^ (**Fig. 6b**) shows that iPSC-derived neurons are relevant for multiple brain-related traits that are associated with the cortex/frontal cortex. Additionally, they provide increased specificity over the tissue-level in many cases, as illustrated for bipolar disorder^38^ with enrichment in both excitatory and inhibitory neuronal subtypes. We further captured certain traits missed in GTEx tissue such as risk tolerance in deep layer excitatory and MGE-like inhibitory neurons, as predicted by Karlsson Linnér, *et al.*^39^. Similarly, we found a strong association between depression and excitatory deep layer neurons, as well as maturing excitatory neurons^40^. We also observed a high enrichment of educational attainment-associated genes in InMGE neurons, concordant with reports of the role of inhibitory neurons in learning and memory^28^. These findings highlight the importance of generating both excitatory and inhibitory neuronal cells within our *in vitro* model.

**Figure 6:**
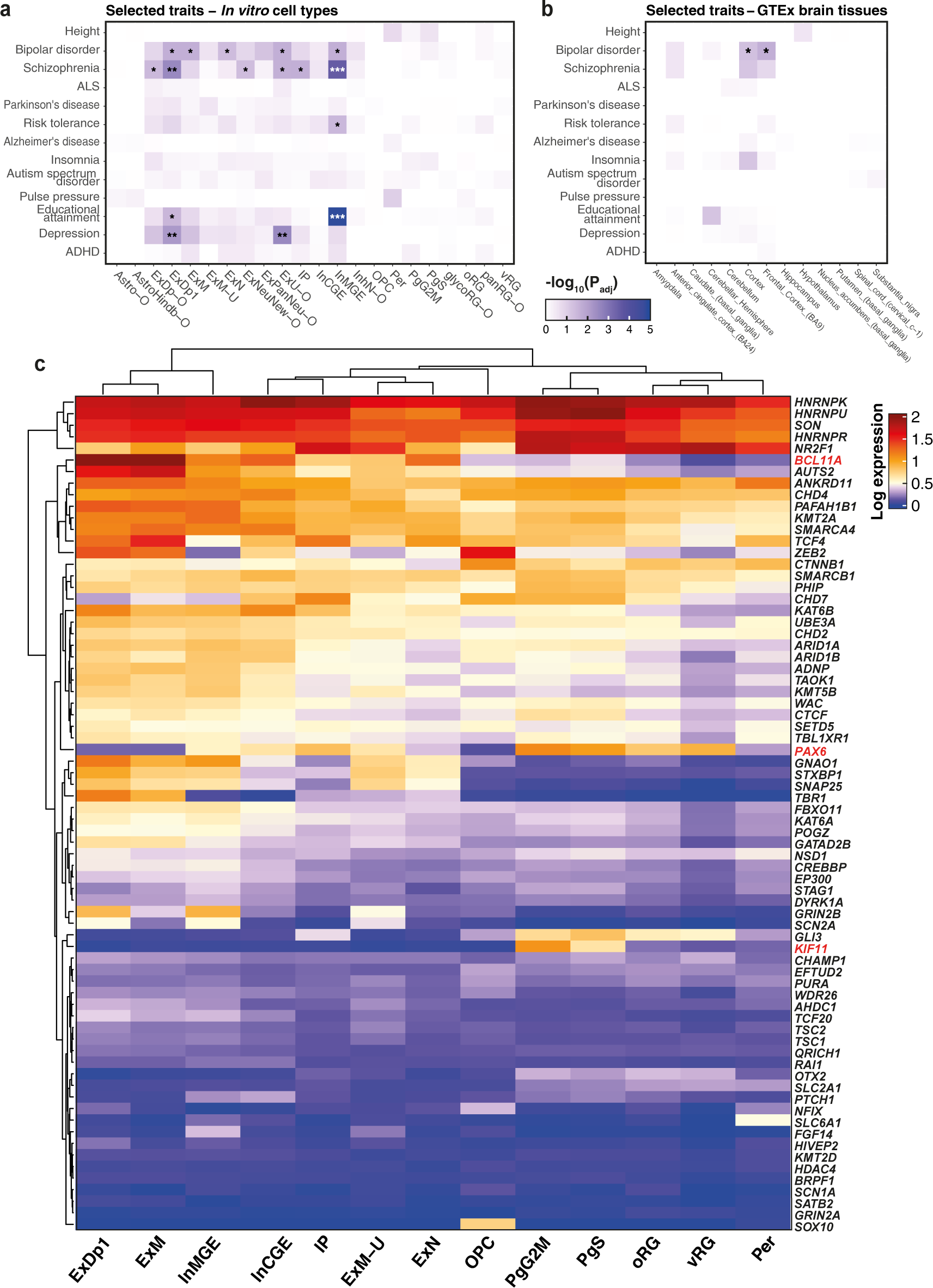
***In vitro* neurons capture brain-relevant traits. (a)** Stratified LD score regression analysis shown for selected brain-related er cell type from our *in vitro* differentiation. Tile color represents the corresponding p values after multiple testing correction across nd cell types, with the significance level indicated as follows: pAdj*<*0.05(*), pAdj*<*0.01(**) and pAdj*<*0.001 (***). **(b)** Stratified re regression analysis as in (a) per GTEx Brain tissue type. **(c)** Heatmap of normalized expression, pseudobulked per cell type, -specific genes associated with developmental disorders (DDD study). Only DDD genes known to be highly intolerant to loss of mutations are illustrated (n=72). Genes highlighted in red represent examples of cell type-specific expression.

In addition to brain-related complex traits, iPSC-derived neuronal models are widely used to model rare NDDs, given their ability to recapitulate cell lineages from very early developmental time points and their transcriptional similarity to fetal, rather than adult cell types. To identify which of our cell types expressed genes associated with NDDs, we evaluated the cell type-specific expression on brain-specific developmental disorder genes from the Deciphering Developmental Disorders (DDD) study^41^. Interestingly, we observed that while approximately 45% of all high- confidence, brain-related DDD genes (n=719) exhibit their highest expression levels in mature neuronal cell types, progenitor cell types also captured another ∼30% of genes associated with developmental delay (**Fig. S7c**). Focusing on a subset of DDD genes (n=72) known for their high intolerance to loss-of-function mutations (**Methods**) highlighted the presence of both neuron- specific (e.g., *BCL11A*) and progenitor-specific genes (*PAX6* and *KIF11*) (**Fig. 6c**). This underscores the importance of using temporal models of neurodevelopment in the context of disease research.

## Discussion

We have performed a comprehensive characterization of iPSC-derived cortical neurons from a widely-used protocol where we combined image-based readouts of cellular morphology with single cell transcriptomics. We describe a proof-of-concept approach, where we applied Cell Painting to a heterogeneous, dynamic system - cortical neurodevelopment - and explored how the joint analysis of imaging and gene expression-based profiles of single cells can contribute to identifying new cellular phenotypes. While scRNA-seq alone allowed us to classify a diverse array of cell types produced, the dynamic nature of developing neurons involves alterations beyond the transcriptome such as in cellular size, shape, and complexity. In neuronal cell types, CP has previously been applied to iPSC-derived neuronal progenitors^42^ as a readout for drug treatment, rather than as a characterization approach. Application of CP in our study throughout the developmental trajectory offered novel insights on the dynamics of morphological readouts, as observed e.g. with mitochondrial channel intensity (**Fig. 3b-c**).

Previous studies have attempted to link bulk gene expression and cell morphology assuming shared biological information between the two, and have found that changes in image-based features are associated with the expression of a subset of genes, often related to the cellular components and organelles stained in the assay^9,15^. The generalizability of the CP assay^43^ means that such direct links of gene expression to cellular components, originally observed in a human osteosarcoma cell line U2OS, are likely reproducible in different cell types. This hypothesis led us to attempt linking these two modalities in our neuronal system. While our sample size is small and limits our ability to generalize our findings beyond our specific set of differentiations, donor- specific variation and the large number of individual cells in our dataset boosted the capacity of the predictive model to link variation in gene expression to CP features. For example, we identified functional terms linked to CP features that recapitulate known or expected biology, such as those linked to ER, supporting our model’s performance. However, beyond organelles, morphology-to- gene expression links are likely to be cell type-specific, making it difficult to compare findings from our study to the previous body of work in non-neuronal cells. To enhance the interpretability of novel associations between image features and their potential biological function, we applied a linear model rather than a machine learning approach, as in previous cross-modality CP-based approaches^44^. In the future, new techniques that allow deriving CP readouts and transcriptomes from the same individual cell would offer a ground truth for model validation.

In general, a multi-modal perspective of any system offers insights that may not be visible from any one assay alone^45^. Here, CP revealed donor-specific ER changes corresponding to transcriptomic differences in ribosomal genes of inhibitory versus excitatory neurons. Although these two cell types transcriptionally cluster together, the CP cluster marked by the lowest ribosomal association, which likely contains low-ribosome inhibitory neurons, groups amongst progenitors. We thus hypothesize that these observed ribosomal/ER-linked differences could reflect differences in maturity between excitatory and inhibitory neurons. Understanding how interneuron production is altered at the level of both gene expression and cellular processes is key to uncover the mechanisms implicated in NDDs, as suggested by recent findings implicating the ER and cytoskeleton in interneuron development and migration^29^. Further, the production of inhibitory cell types in our *in vitro* system highlights the importance of a human-specific model to recapitulate fetal development, since inhibitory neurons are not known to be produced in the cortex of other model organisms such as mice^24^.

A common challenge in modeling neurodevelopment using pluripotent stem cells, regardless of the applied protocol, lies in determining whether the cell types generated *in vitro* accurately mirror the transcriptional signatures and developmental trajectories observed *in vivo*. Previous studies in organoids have linked *in vitro*-specific cell states to aberrant oxidative or glycolytic stress^4,34^. These alterations in energy-associated pathways are frequently associated with the limitations of the culture media to provide essential nutrients to cells, especially within the necrotic cores of organoids^34^. In a recent study^35^, transcriptomic differences between cells from human neural organoids (comprising 26 protocols) and developing human brain (first-trimester)^46^ were associated with the upregulation of canonical glycolysis and mitochondrial ATP synthesis-coupled electron transport in organoids. Notably, canonical glycolysis was adopted as a proxy for cell stress given its association with expression differences observed between organoid and primary cells. Similarly, we observed that low-quality cells in our 2D culture system showed distinct metabolic and energetic states compared to high-quality cells of the same type. For instance, pathway activation scores in glycolysis differed among progenitors, and in steroids biosynthesis among excitatory neurons. A fraction of these low-quality cells could indeed be linked to an identity exclusively seen in organoids, most of them being either pan-neuronal or pan-radial glial. Mis-annotating these low-quality cells could potentially bias downstream findings in disease models if not accounted for.

Overall, based on our observations, single cell transcriptomics remains a far more in-depth tool for cell type characterization than CP. Indeed, only larger supertypes of cells (progenitors versus neurons) were resolved with the standard CP assay, potentially due to the generic nature of the included dyes. Replacing generic dyes with those specifically targeting neurons could improve cell type granularity^16^, as could increasing image magnification for enhanced resolution of individual organelles. This could help us to better understand low quality cell types within CP data, currently not addressed in our study due to the model limitations. Further, to capture morphological heterogeneity, we analyzed CP readouts at a single cell level instead of averaging CP features by well after cell-segmentation as is often done, which came at the cost of increased noise^13^. Although lower seeding density in culture may improve the accuracy of segmentation, we have previously observed that the viability of developing neurons is compromised in sparser culture conditions. It is also uncertain whether the 384-well format of the CP assay, as compared to 35mm dishes used for scRNA-seq, impacts cell type production.

Finally, we applied stratified LD score regression analysis to show that our *in vitro* model is relevant to several brain-related traits associated with the human cortex, capturing more trait heritability than GTEx brain tissues^37^. This is expected due to the better resolution with cell type-specific expression, but nonetheless illustrates the power of being able to generate and study the most disease-relevant cell types *in vitro*. In our results, InMGE neurons stand out by capturing the highest number of traits (n=4), with educational attainment being the most significant association. This underscores the vital role of interneurons in maintaining the balance between excitation and inhibition in neural circuits during development, with perturbations of this system having potential implications for processes like learning and memory^28^. Furthermore, we observed distinct temporal and cell-type specific expression of genes associated with developmental disorders in the brain, emphasizing the need to incorporate temporal models in neurodevelopment for disease modeling.

In conclusion, we have performed an in-depth characterization of a widely used and highly disease-relevant cortical model using joint profiling of cell morphology and gene expression. We provide a novel framework for combining image-based phenotypes with transcriptomics in single cells and highlight the potential of cell morphology measurements to capture dynamic cell states and biological processes that are complementary to those captured by scRNA-seq. We expect this approach to be widely useful in disease modeling^10,42^ and other functional genomics studies in diverse *in vitro* neuronal systems.

## Materials and Methods

### Human iPSC culture

The human iPSC lines HEL11.4, HEL47.2, HEL61.1, HEL61.2, HEL62.4 and HEL82.6 used in this study were acquired from the Biomedicum Stem Cell Centre (University of Helsinki, Finland) (**Table 1, S1**). The cells were grown on vitronectin in Essential 8 and Essential 8 Flex media (**Table S5**) at 37°C/5% CO2. iPSC maintenance in culture was performed according to HipSci guidelines: https://www.culturecollections.org.uk/media/109442/general-guidelines-for-handling-hipsci-ipscs.pdf). Cells were clump-passaged in ratios ranging from 1:4 to 1:8 using 0.5 mM EDTA diluted in DPBS-/-. Y-27632 (10 µM) was used for better cell survival at thawing. Cells were tested for mycoplasma with the MycoAlert kit and all lines tested negative after thawing both during iPS cell culture and neural differentiation.

**Table 1.** Cell lines used in the study with their corresponding donor information, derivation method, registration (https://hpscreg.eu/) and karyotype cytogenetic nomenclature.

| Name | Cell Type | Diagnosis | Sex | Derivation method | Characterized as iPSC | Marker expression | Ref | Registry info | Karyotype |
| --- | --- | --- | --- | --- | --- | --- | --- | --- | --- |
| HEL11.4 | Skin fibroblast | Unaffected | Male | Retrovirus | Yes | Yes | <sup>59</sup> | UHi007-B | 46,X,inv(Y)(p11q11),add(1)(q12q21) |
| HEL47.2 | Skin fibroblast | Unaffected | Male | Sendai virus | Yes | Yes | <sup>60</sup> | UHi007-A | 46,X,inv(Y)(p11q11) |
| HEL62.4 | Skin fibroblast | Unaffected | Female | Sendai virus | Yes | Yes |  | UHi020-A | 47,XX,+12[2]/46,XX[18] |
| HEL61.1 | Skin fibroblast | Unaffected | Female | Sendai virus | Yes | Yes |  | UHi021-A | 46, XX |
| HEL61.2 | Skin fibroblast | Unaffected | Female | Sendai virus | Yes | Yes |  | UHi021-B | 46, XX |
| HEL82.6 | Skin fibroblast | Unaffected | Female | Sendai virus | Yes | Yes |  | UHi022-A | 46, XX |

### Cortical neuron differentiation

The iPSC lines were differentiated into cortical progenitors and neurons using an established differentiation protocol^11^ with minor modifications (hereafter referred to as ‘original’) and a modified version from day 11 post-induction (**Table S2**).

For neural inductions, 2-3 80% confluent plates of iPS cells were detached using 0.5 mM EDTA and were plated on a Matrigel plate (1:100 in DMEM/F12) in E8 medium supplemented with Y- 27632 (10 µM). Dual-SMAD inhibition was initiated the following day using Neural Maintenance Medium (NMM) supplemented with SB431542 (10 µM) and LDN-193189 (200 nM) (**Table S5**). On day 11 post-induction, cells were split in clumps in 1:2 ratio onto laminin-coated dishes using either mechanical dissociation by gentle scraping (modified) or using Dispase (original). The following day, the media was changed to NMM containing bFGF (0.2 µg/ml) for four days. After expansion, cells were split (1:2) with EDTA only in the modified protocol. Alternatively, a few replicates (techRep column, **Table S2**) for days 40 and 70 were dissociated with Dispase as per the original protocol.

At day 17, the plates to be assayed at day 20 (modified protocol) were split 1:2 as small clumps (using 0.5 μM EDTA) onto laminin-coated plates for scRNA-seq analysis. Additionally, cells were plated in 1:6 ratio on 24-well plates with coverslips for immunocytochemistry and in 1:120 ratio on 384-well plates for Cell Painting. Cells assayed at days 40 and 70 were split as described in **Table S2**.

All cells were frozen down at day 28 or 29 and thawed as per the original protocol. Cells were frozen down in 1:1 ratio and were plated onto laminin-coated plates at thawing. Final plating for days 40 and 70 was done at day 35 when cells were passaged with Accutase. Cells were plated for scRNA-seq (1.5 million cells/35mm dish), immunocytochemistry (75-300,000 cells/24-well plate) and Cell Painting (5,000 cells/384-well plate) onto poly-L-ornithine and laminin-coated plates. Poly-L-ornithine solution was diluted to 0.01% with sterile water, coated overnight at +4°C after which the wells were washed 3 times with sterile water. Laminin was diluted in DPBS-/- and the plates were incubated at 37°C for 4h.

scRNA-seq samples profiled at days 20, 40 and 70 were not taken from the same continuous differentiation. Days 40 and 70 were sampled from one round of differentiation, containing 4 donors (HEL61.2, HEL11.4, HEL62.4, HEL82.6) and with technical replicates as specified in **Table S3**. An additional round of differentiation was run to profile Cell Painting samples in all three time points, and in addition, day 20 scRNA-seq samples were obtained from this batch. These samples were only differentiated using the modified version of the protocol. This time point incorporated two additional iPSC lines, HEL61.1 (clone of HEL61.2) and HEL47.2 (derived from the same donor as HEL11.4, but generated from different parental fibroblasts), requiring the barcoding technology of CellPlex to demultiplex donor identity. For this time point, two independent inductions were replicated one week apart (batchRep column, **Table S3**).

### Cell preparation for single cell RNA sequencing

Developing cortical neurons were analyzed for experiments on days 20, 40 and 70 post neural induction. The cells were prepared for scRNA-seq as follows: The wells were washed up to three times with DPBS-/- after which they were incubated in Accutase for 5 min at 37°C. Cells were dissociated into a single cell suspension by pipetting and added into 5 ml of 0.04% BSA in DPBS-/-. Cells were centrifuged at 180 RCF for 5 min and supernatant was removed. Cells were resuspended in 0.04% BSA and centrifuged twice more. Final resuspension of cells was done in 100 µl of 0.04% BSA after which the cells were filtered through 40 µm FlowMe filters, followed by counting and estimation of the cell viability using Trypan Blue.

### Single-cell RNA-sequencing library chemistry and sequencing

Single-cell gene expression was profiled from the three time points using 10x Genomics Chromium Single Cell 3’ Gene Expression technology. Only at day 20, Cell Multiplexing technology platform (3’ CellPlex Kit) was used to demultiplex the identity of clonal cell lines. For all the time points, 10x libraries were generated using the Chromium Next GEM Single Cell 3’ Gene Expression version 3.1 Dual Index chemistry. The sample libraries were sequenced on Illumina NovaSeq 6000 system using read lengths: 28bp (Read 1), 10bp (i7 Index), 10bp (i5 Index) and 90bp (Read 2) (**Table S4**).

### Genotyping

To allow for donor demultiplexing during downstream analysis, iPSC lines were genotyped using SNP arrays. For this, cells were pelleted in DPBS-/- and DNA was extracted using Nucleospin DNA columns. Genotyping was performed on Illumina Global Screening Array with added GSAFIN SNPs specific for the Finnish population.

### Immunocytochemistry and image acquisition

Cells were washed three times with DPBS+/+, fixed with 4% paraformaldehyde for 15 min followed by three DPBS washes. Cells were then permeabilized in 0.2% Triton X-100/DPBS for 15 min at RT (**Table S5**). Coverslips were washed three times in PBST (0.1% Tween-20) followed by blocking at RT with 5% BSA/PBST for two hours. Cells were incubated in primary antibodies in 5% BSA/PBST overnight at 4°C (**Table S6**). Following overnight incubation, coverslips were washed with PBST for 15 min three times, followed by incubation in secondary antibodies in 5% BSA/PBST for one hour. The cells were finally washed three times with DPBS for 10 min and coverslips were plated on glass slides with mounting media containing DAPI. Fixation for lines HEL62.4 and HEL82.6 was performed on day 55 rather than day 70 due to neuron detachment from the coverslips.

Imaging for day 20 ICC was performed using a Zeiss Axio Observer.Z1. The objective used was a Plan-Apochromat NA 0.8 at 20x magnification. Samples were imaged with HXP 120V light source with the 45 Texas Red, 38HE GFP and 49 DAPI wavelength fluorescence filters. Images were acquired using an Axiocam 506. For days 40 and 70, imaging was performed using a Zeiss Axio Imager 1 with the same objective and light source. Fluorescence filters used were 64HE mPLum, 38HE GFP and 49 DAPI. Images were acquired using a Hamamatsu Orca Flash 4.0 LT B&W.

### Cell Painting assay and image acquisition

Cells were phenotyped using Phenovue Cell Painting Kit for 384-well plates following kit guidelines based on Bray *et al.* (2016)^7^. Cells were plated on day 17 (for day 20) or day 35 (for days 40 and 70) on PhenoPlate 384-well microplates and incubated in 37°C at 5% CO2 until assay time points at days 20, 40 and 70. Staining solution 1 was added for live labeling of mitochondria (PhenoVue 641 Mitochondrial Stain) and incubated in the dark for 30 min at 37°C. Cells were fixed with 3.2% PFA for 20 min at room temperature, and then were washed and incubated with 0.1% Triton X-100 followed by HBSS washes. Finally, staining solution 2 was added to the cells to label nuclei (PhenoVue Hoechst 33342 Nuclear Stain), ER (PhenoVue Fluor 488 - Concanavalin A), Golgi apparatus (PhenoVue Fluor 555 - WGA), nucleic acid (PhenoVue 512 Nucleic Acid Stain) and cytoskeleton (PhenoVue Fluor 568 - Phalloidin) and washed again with HBSS prior to imaging.

The cells were imaged with PerkinElmer Opera Phenix High Content Screening System using the Harmony Software v4.9. The imaging was done using the 40x NA 1.1 water immersion object and with the following lasers: 405 nm (emission window 435-480 nm), 488 nm (500-550 nm), 561 nm (570-630 nm), and 640 nm (650-760 nm), resulting in Golgi apparatus, nucleic acid and cytoskeletal dyes being captured in the same channel (CGAN). Images were captured using the Andor Zyla sCMOS camera (2160 x 2160 pixels; 6.5 μm pixel size). For each well, 28 fields on 3 planes were acquired. Amongst images taken from multiple planes (n=3), the z-stack with the highest intensity per channel was selected for analysis.

### scRNA-seq data pre-processing and dimensional reduction

Raw data processing and analysis were performed using 10x Genomics Cell Ranger v6.1.2 pipelines “cellranger mkfastq” to produce FASTQ files and “cellranger multi” to perform alignment, filtering and UMI counting. mkfastq was run using the Illumina bcl2fastq v2.2.0 and alignment was done against human genome GRCh38. Day 20 samples were demultiplexed based on multiplexing barcode sequences using the cellranger multi pipeline. Cell recovery was 22,628 and 18,486 from the two 10x samples sequenced, with 50.89% and 73.61% cells assigned to a cell line (singlet rate), respectively, resulting in approximately 4,000 cells captured per cell line across technical replicates (**Table S4**). In 10x samples from days 40 (n=3) and 70 (n=2), donors were pooled at the time of sequencing. We assigned donor identity to each cell with *demuxlet*^47^ by leveraging common genetic variation from the same donors previously genotyped (see Methods, Genotyping section). *Demuxlet* was run using a default prior doublet rate of 0.5. We only retained those (singleton) cells that could unambiguously be linked to a donor (average of 69.5% per pool), resulting in 5,011 cells on average per donor at each timepoint.

scRNA-seq data was analyzed using the *Seurat* R package v4.1.1 (Stuart et al., 2019) using R v4.1.3. To exclude low-quality cells from analysis, we discarded cells with either less than 2,000 genes expressed or more than 8,000, as well cells presenting more than 15% of reads mapping to the mitochondrial DNA. Additionally, it was ensured that each 10x sample contained between 10-20% reads mapping to ribosomal protein transcripts on average, as expected from neuronal populations. Following filtering, the 10x samples from all three timepoints were merged and genes expressed in <0.1% of cells in the merged dataset were removed.

Gene expression counts were normalized by total expression, with a default scale factor of 10,000, and then transformed to log-space. Then, this matrix of log-normalized counts was scaled while regressing out cell cycle scores^48^, depicted as the difference between G2/M and S phase scores to preserve inherent differences between cycling and non-cycling cells. Dimensionality reduction was performed via Principal Component Analysis (PCA) using previously identified highly variable genes (n=3,000). *Harmony* (v.0.1.0) was used to batch-correct the PCA embeddings from all 10x samples^49^. Based on the top 15 batch-corrected PCs, we constructed a KNN graph and clustered the cells with a resolution of 0.8, using Seurat functions *FindNeighbors()* and *FindClusters()*, respectively. We then generated a UMAP embedding again using the top 15 batch-corrected PCs from *Harmony*.

### scRNA-seq cell type annotations from *in vivo* fetal brain

The primary reference dataset used for cell type annotation was the publicly available human fetal scRNA-seq data from Polioudakis et al.^23^, obtained from the CoDEx online interface (http://solo.bmap.ucla.edu/shiny/webapp/). It was selected given the data and code availability, plus the metadata with regional specificity of the developing neocortex. The raw matrix of counts was log-normalized to a scale factor of 10,000 counts and we identified the top 3,000 highly variable genes. Based on the expression of G2/M and S phase markers, we calculated cell cycle phase scores. Then, counts were scaled, regressing out ‘Number_UMI’, ‘Library’ and ‘Donor’ and the difference between G2/M and S cell cycle scores. Dimensionality reduction was performed via PCA on the top 3,000 highly variable genes. We then batch-corrected the PCA embeddings with *Harmony* specifying the library as a covariate and used the harmonized dimensional reduction as an embedding to project our *in vitro* dataset. To transfer the cell type labels from the fetal reference to our *in vitro* query dataset, we used a two-step anchor-based approach implemented in Seurat: first running ‘FindTransferAnchors’ using the first 30 batch-corrected PCA from the reference embedding, and then “MapQuery”. Further, the tag of ‘Unmapped’ was assigned to cells that did not achieve a mapping score of >0.5 for any cell type label. Correlation analysis of the annotated cell types between the *in vitro* and reference datasets was performed similar to Bhaduri *et al.* (2020)^4^. On-diagonal and off-diagonal means of Pearson correlation coefficient were calculated. We classified cells in bins of low and high quality within a cell type assigned based on their best mapping score. Those cells with a score>0.5 were referred to as high quality for a predicted cell type, while cells with a score<0.5 were considered as low-quality cells. Additionally, cell types with less than 50 cells were not considered for any downstream per-cell type analyses (in this case, only ExDp2).

### Pseudotime analysis

The pseudotime trajectory was constructed using R package *monocle3* (v1.2.9). The Seurat object was converted to the *monocle3*-compatible cds object type using the function ‘*as.cell_data_set()*’ followed by pre-processing with 100 dims, alignment by 10x sample and clustering at a resolution of 1e-4. The ‘*learn_graph()*’ function was used with default parameters to construct the trajectory and the cells were ordered along the trajectory using a principal node rooted in the time point day 20. Genes that vary across the trajectory were identified using ‘*graph_test()*’ by setting the argument ‘*neighbor_graph’’* to *”principal_graph*”. The resulting genes were grouped into modules based on ‘*find_gene_modules()*’ after evaluating modularity using different resolution parameters {10^-6^, 10^-5^, 10^-4^, 10^-3^, 10^-2^, 10^-1^}.. Finally, the expression of these modules was aggregated per cell type using the function ‘*aggregate_gene_expression()*’.

### GO enrichment

Overrepresentation analysis using gene ontology (GO)^50^ was performed using *clusterProfiler* v4.2.2 and *org.Hs.eg.db* v3.14.0. The gene universe consists of 23,289 genes present after filtering out those expressed in <0.1% of cells (see Methods, scRNA-data pre-processing), and mapped to ENTREZ and ENSEMBL gene IDs. The *enrichGO* function from *clusterProfiler* was used to find enriched terms of all categories (‘BP’, ‘CC’ and ‘MF’) per previously identified gene module and top 15 terms per analysis (passing a qvalueCutoff=0.05) were visualized using *enrichplot* v1.14.2.

### GSVA

Gene expression was aggregated per cell type based on mean log-normalized expression values to generate pseudobulked data. Gene set variation analysis was performed on pseudobulked cell types using the R package *GSVA* v1.49.4 and gene sets obtained from the Molecular Signatures Database (MSigDB)^51^ compiled in the R package *msigdbr* v7.5.1. Selected pathways from KEGG (**Fig. 3d & 5c**) or Gene Ontology: Cellular Components (GO CC) containing the ‘MITOCHONDRIAL’ term (**Fig. 3e**) were tested for enrichment across cell types using the function ‘gsva’ with a min.sz filter of 15 and max.sz of 500.

### Organoid mapping

Reference scRNA-seq data from cortical organoids was obtained from Bhaduri et al.^4^, via the UCSC Cell Browser (https://cells.ucsc.edu/?ds=organoidreportcard). As in the original publication, the raw count matrix was pre-processed following original filtering steps to remove cells with fewer than 500 genes expressed or with an excessive mitochondrial count fraction (>10%). Then, gene counts were normalized to a scale factor of 10,000 counts and natural-log transformed and 3,000 highly variable genes were computed. Based on the expression of G2/M and S phase markers, we calculated cell cycle scores, and the difference between G2/M and S was regressed out during data scaling. We performed PCA dimensionality reduction using the top 3,000 highly variable genes. We used the first 30 PCA to project the organoid reference to our in vitro dataset as described for fetal mapping. This reference mapping was the second step in producing the final cell type annotation, as we only assigned the organoid cell type labels to those cells that were classified as ‘unmapped’ from the fetal reference-based (first-step annotation). Those cells that did not achieve a maximum mapping score of >0.5 for any cell type label in any of the two mappings were finally tagged as “Unmapped”.

### Differential abundance testing

We tested for differential abundance between the cell types produced by the two versions of the protocol (original vs modified) using *miloR* v1.4.0^36^. This analysis was performed solely with samples from day 40 due to the representation of all donors in each protocol version (**Table S2**). The KNN graph was constructed using the *buildGraph()* function with 15 dimensions (d=15) and 30 nearest-neighbors (k=30), followed by *makeNhoods()* using the same number of dimensions and sampling 20% of the graph vertices (prop=0.2). In both cases, the dimensionality reduction from Harmony after batch-correcting the PCA was used. To calculate the distance between neighborhoods, we used the function *calcNhoodDistance()* with 15 dimensions (d=15) from Harmony. The neighborhoods were tested for differential abundance between protocols by running *testNhoods()* with the design:

∼ donor + protocol

The resulting differentially abundant neighborhoods were annotated with cell types from the two-step annotation using *annotateNhoods()*, and neighborhoods that were not homogeneously composed of a single cell type (fraction of cells from any given cell type <0.7) were annotated as ‘Mixed’. The differential abundance fold changes were visualized using a beeswarm plot with default significance level for Spatial FDR (<0.1).

### Gene Set Enrichment Analysis (GSEA)

GSEA was performed on differentially expressed genes between pan-neuronal and inhibitory cell types (**Fig. 4d**) or between excitatory and inhibitory cell types (**Fig. 5b, S5c**) using *fgsea* v1.20.0. The function ‘fgseaMultilevel’ was used on the ranked list of DEGs using KEGG gene sets from msigdbr and filtering for minsize=10, maxsize=500, eps=0 and nPErmSimple=10000. Results were ordered by NES and filtered for padj<0.05 before visualizing the top (maximum) 20 results per analysis. Additionally, in sections 4 & 5, the genes driving specific pathways were identified based on the ‘leadingEdge’ of each pathway and represent the core of the gene set enrichment’s signal^18,19^. The module score for each of these core sets of genes was computed per single cell using ‘AddModuleScore’ from *Seurat* with default parameters.

### Cell Painting image processing

The acquired images were processed using *CellProfiler* 4.2.1^52^. The workflow file is available at Github (CellPainting/1_primary_analysis/cellProfilerWorkflow.cppipe). Briefly, quality metrics such as blurriness and saturation were measured for each image. Nuclei were then segmented using the Hoechst channel using the minimal cross-entropy method. Then, an image summing up the channels and excluding Nucleus staining was generated to segmentate the cells (**Supplementary Methods Fig. 1a**). Neurites were algorithmically enhanced on summed images prior to segmentation. Cell segmentation was done by propagation from the nucleus using the Otsu method. Cytoplasms were identified by subtracting nuclei area to the cell. For each object type (Cell, Nuclei, Cytoplasm), a large number of features were measured per channel. A description of each feature can be found in the CellProfiler manual (https://cellprofiler-manual.s3.amazonaws.com/CellProfiler-4.0.4/help/). Those features were metrics relative to channel intensity, texture or object shape and size. To export images, intensity of each channel was rescaled and attributed to a color. Channels merged to create one image per field in the well. An overlay of the well is then created for each field using a python script (CellPainting/1_primary_analysis/makeComposite_stitch.py).

### Analysis of quantified features of Cell Painting

The feature data frames were analyzed by R 4.2 using the *oob* package (https://github.com/DimitriMeistermann/oob). First, images underwent a filtering process based on the PowerLogLogSlope, a blur metric. For each field, the average PowerLogLogSlope was calculated across the four channels. We retained images with an average value greater than - 2.16, which is determined based on the distribution. Additionally, images identified as blurry by Cell Profiler are excluded. Any objects associated with discarded images are removed from the dataset. Cells exhibiting extreme values for cytoplasm area (≤1000 or ≥100000) or too small nucleus (≤10000) are also eliminated, as they are indicative of poor segmentation. Cells with too low nucleus channel intensity were also excluded (≤0.003), as well as being outlier in a projection with nucleus intensity as x-axis and sum of other channels as y-axis. Outliers were determined by a uniform kernel density estimation with bandwidth=1, and removed if density ≤0.0001.

Features were then regularized to approximately follow a normal distribution. This was done by examining each feature distribution and classifying them. We defined 6 feature distribution families (**Supplementary Methods Fig. 8, Supplementary Methods Table 1**) and applied a specific transformation for each. For example, intensity features underwent a log2(x+1) transformation. Range of each feature was then scaled to [0,1000].

The dataset, originally containing 1,186,458 cells, was reduced for ensuring a relatively balanced contribution from each experimental population (time-point × cell line) and reducing the computation time. The subsampling was aimed to reach a range of 50,000 to 60,000 cells. This specific range was determined using bootstrapping, which involved repeatedly sampling the dataset and assessing the correlation between the subsamples and the original dataset. When 50,000 cells were selected, the average correlation was approximately 0.98. Subsampling was carried out based on experimental populations, considering differentiation day and cell line, with a maximum of 2000 cells drawn if the population exceeded this size.

Feature selection was conducted through a multi-step process. Initially, a feature graph was constructed, and edges were established between features if their correlation exceeded 0.99. Modules were subsequently identified from this graph using the function “cluster_fast_greedy” from the *igraph* package^53^. Within each of these modules, the most parsimonious feature was retained for further analysis. The features associated with spatial measures (x-y locations), Zernike-related values, and Cel_Neighbors_AngleBtNghbors_Adjacent were excluded from further analysis. Their signals did not exhibit a discernible pattern and, as a result, posed a potential risk of introducing noise into subsequent analyses.

Finally, a temporary cell clustering was performed (see next paragraph), and the pictures of cells from each cluster was assessed, revealing two clusters composed of image artifacts or dying cells. Those clusters were removed from downstream analysis. We obtained a feature matrix consisting of 223 features and 54,415 cells.

A UMAP and Leiden clustering analysis was conducted using the *oob* package with a specified parameter of “n_neighbors = 20.” Subsequently, feature modules were identified by performing hierarchical clustering on the features, utilizing a covariance distance matrix. The number of modules was determined using the derivative loss method. This process resulted in the identification of a total of 18 modules, and they were named based on an examination of their content. To determine module activation scores, the first component of a Principal Component Analysis (PCA) was extracted from the matrix, which contained the features of each module for all cells. The web interface for visualizing the CP dataset was coded using the d3.js framework.

### Integration of scRNA-Seq and Cell Painting

For the purpose of the multi-modal analysis, scRNA-seq data was reprocessed to enhance comparability between CP features and gene expression, aiming to align these datasets as closely as possible. The used set of cells is consistent with those used in the primary scRNA-seq analyses (refer to the section on scRNA-seq data pre-processing for more details). The genes with average expression < 0.005 were filtered out and normalization was performed using the computeSumFactors function from *scran*^54^. Batches were corrected using fastMNN with k=5 from the *batchelor* package^55^.

To perform the integration, only the common experimental population (combination of differentiation day and cell line) were selected, then 3 metacells were created per experimental population per modality. The metacells were created by randomly attributing cells from each experimental population to one of three metacell. Feature values or gene expressions were then averaged by metacell. This led to 2 matrices of 42 metacells. These matrices were used to train Lasso regressions models using *sklearn*^56^ from Python (alpha=0.02, max_iter = 10000). For each experimental population, two metacells from each modality were used for the training and one for cross-validation. The intercepts and regression coefficients were then exported to a matrix that was used to predict CP features from gene expression. This matrix was used to build the predicted Cell Painting feature matrix of the scRNA-Seq dataset. CP features markers of each CP cluster were computed in the CP feature matrix, and CP features markers of each cell type in the predicted CP feature matrix. This was done using getMarkers from the *oob* package. Subsequently, two marker score matrices were generated and correlated to obtain **Fig. 2e**.

Parallelly to the lasso regression models, regular linear regressions were computed with the same formula (*CP feature ∼ genes*) with the aim to provide one value per gene for each CP feature. This enabled the use of GSEA to enrich CP features, using the regression coefficients as GSEA input scores. Prior to the enrichment, a median of coefficients was computed per CP module to perform the enrichment per module with KEGG and GO databases as gene set databases.

For each cell from the scRNA-Seq dataset, the predicted CP feature matrix was used along the Seurat cell cycle scores annotation to build two lasso regression models: one to predict S.score, the other to predict the G2M score with a formula on the form of *score ∼ CP features*. The models were then used to predict the cell cycle scores values in the CP dataset.

### AMI

Adjusted Mutual Information (AMI) was computed using the aricode package for R^57^.

### Stratified linkage disequilibrium (LD) score regression analysis

Positive markers per annotated cell type from our *in vitro* dataset were determined using *Seurat FindMarkers()* function. A 100 kb window was added on either side of each of these genes using the GenBank reference genome version NCBI:GCA_000001405.14 from GRCh37.p13 (https://ftp.ensembl.org/pub/grch37/current/fasta/homo_sapiens/pep/) and LD scores computed. We then investigated heritability enrichment for 79 traits (**Table S7**) given our cell-type specific annotations (n=22 cell types) using stratified LD score regression implemented in LDSC (*LDSC* v1.0.0)^14^, with the full set of genes expressed in at least <0.1% of cells (n=23,289) as the control gene set. As a comparison, we ran the same analysis using annotations for 13 brain tissues from GTEx^37^. Multiple testing correction was performed across resulting trait-specific p-values across all cell/tissue types using the Benjamini-Hochberg Procedure.

### Expression of brain-related developmental delay genes

Developmental delay associated genes from the DDD study were obtained from https://www.ebi.ac.uk/gene2phenotype (version 28_7_2023). The geneset was filtered for loss of function (absent gene product) brain-specific genes of ‘definitive’ confidence and with autosomal monoallelic requirement. Finally, genes were filtered for loss-of-function observed/expected upper bound fraction (LOEUF) score <0.3 in order to limit the analysis to genes with higher probability of large consequence on cellular phenotypes. LOEUF information was downloaded from gnomAD v4^58^. The selected genes’ scaled expression was plotted per pseudobulked cell type from the fetal annotation.

## Data availability

Raw scRNA-seq data and genotype information from the cell lines used in the *in vitro* cortical differentiation will be made available on the European Genome-Phenome Archive. Additionally, raw image data from CP are deposited in the EMBL-EBI BioImage Archive under accession number S-BIAD969. Processed Cell Painting data (e.g. feature matrix, model coefficients, cell annotations) are available from Zenodo (https://zenodo.org/doi/10.5281/zenodo.10213789).

## Code availability

Code used to analyze the scRNA-seq and Cell Painting datasets is available at https://github.com/Kilpinen-group/cortical_diff_code/. Additionally, an online interface linking the CP UMAP to individual images to facilitate exploration of the CP dataset is available at https://vm3725.kaj.pouta.csc.fi/.

## Acknowledgements

This work was primarily funded by the Helsinki Institute of Life Science (HiLIFE), University of Helsinki (UH). We also acknowledge additional financial support from the Doctoral Programme in Integrative Life Science, UH (AS) Maud Kuistila Memorial Foundation (RW), and the Sigrid Juselius Foundation (HK, PPC). The cell lines used in this study were obtained from the Biomedicum Stem Cell Center (BSCC), supported by HiLIFE and Biocenter Finland. Genotyping was performed at the Institute for Molecular Medicine Finland (FIMM) Genomics unit, supported by HiLIFE and Biocenter Finland. The authors would like to thank the FIMM Single-Cell Analytics unit, supported by HiLIFE and Biocenter Finland, for single-cell RNA-sequencing services, and Dr. Lassi Paavolainen and the FIMM High Content Imaging and Analysis unit for assistance with Cell Painting and for imaging and image data analysis. Imaging for ICC was performed at the Biomedicum Imaging Unit, Helsinki University, Helsinki, Finland, with the support of Biocenter Finland. The authors also wish to acknowledge CSC – IT Center for Science, Finland, and the Institute of Molecular Medicine Technology Center for computational resources, and Dr. Lewis Evans for providing feedback on the manuscript.

## Author Contributions

A.S, R.L and R.W conducted the culture of the chosen iPSC lines, ran the differentiation process of cortical neurons, and prepared the cells for single-cell data sequencing. They also conducted the cell profiling for immunochemistry assays, processed cells for Cell Painting, and performed image acquisition. A.S. and P.P. processed single-cell data and performed the reference mapping for cell type annotation. A.S. and D.M. led the main data analysis and interpretation: A.S. drove the single-cell RNA-seq analysis (differential expression, pseudotime and enrichment analysis), while D.M. conducted Cell Painting image processing, analysis, and integration with single-cell data. Z.Y. run the stratified LD score regression and A.G provided the GWAS summary statistics for brain-related traits. H.K., P.P. conceived the study and provided guidance. A.S., D.M. and H.K wrote and edited the manuscript, as well as produced all the figures.

## Declaration of interests

The authors declare no competing interests.

## Notes

### Competing Interest Statement

The authors have declared no competing interest.

### Summary of Updates

Textual restructuring and minor edits throughout text; restructuring of figure panels in figs 1 and 2; figures 4 and 5 interchanged.

