## Supplementary Methods Fig for "Joint profiling of cell morphology and gene expression during *in vitro* neurodevelopment"

### Supplementary Methods Figures

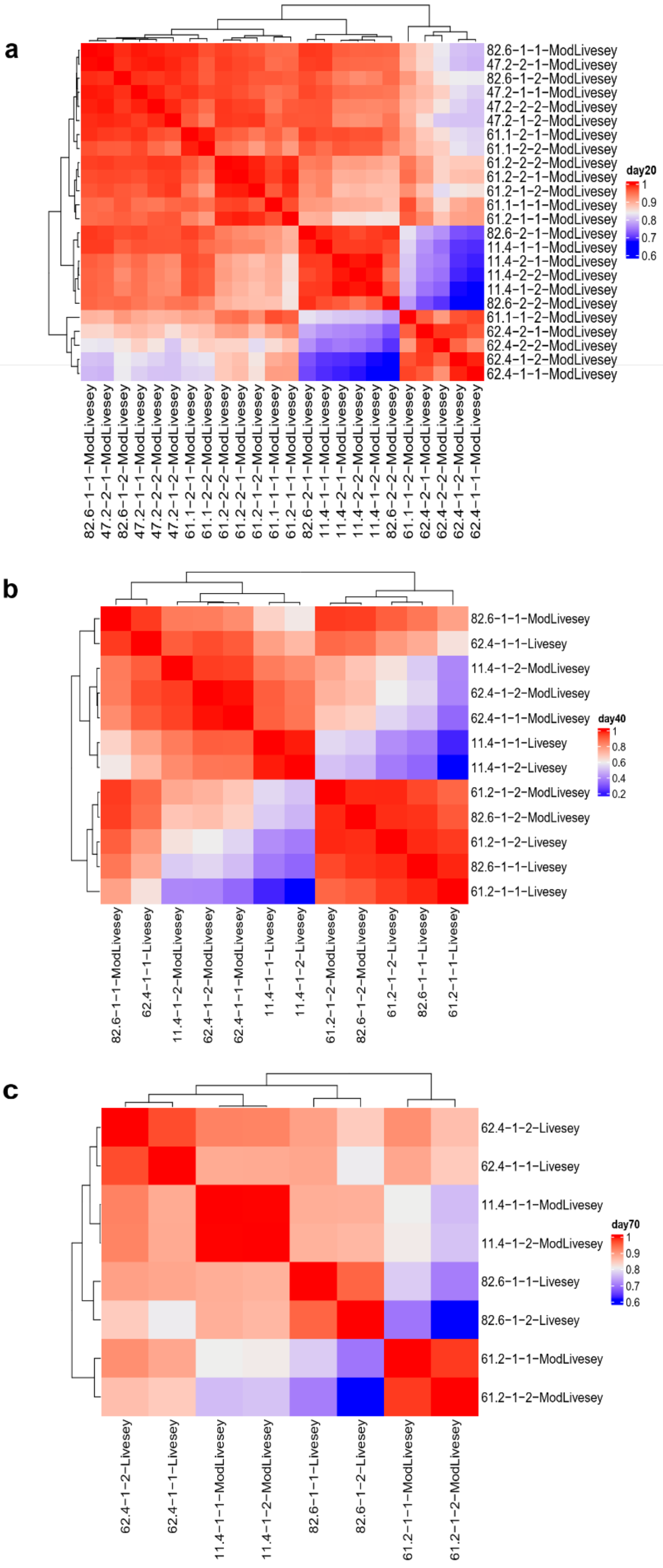


Supplementary Methods Fig. 1

**Supplementary Methods Fig. 1. scRNA-seq quality control. (a)** Pearson correlation of cell type composition between differentiation replicates at day 20. Sample labeling is donor-batchRep-techRep-protocol with numbering corresponding to **Table S3**. **(b-c)** Pearson correlation as in (a) for samples at day 40 **(b)** and day 70 **(c).**

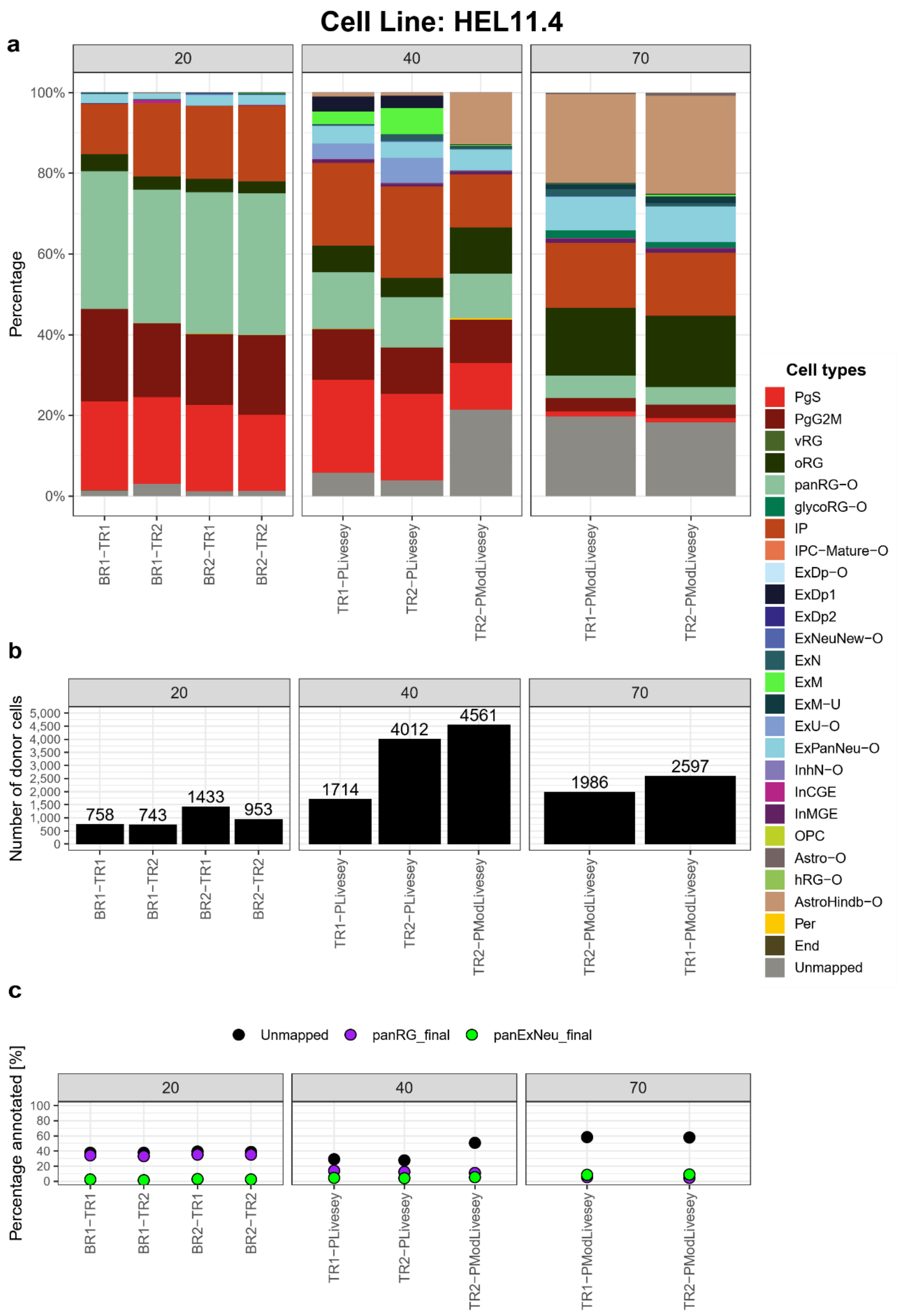


Supplementary Methods Fig. 2

**Supplementary Methods Fig. 2. Reproducibility of replicates in scRNA-seq: HEL11.4. (a)** Cell type composition per replicate, per differentiation time point within cells from cell line HEL11.4 (BR - batch replicate, TR - technical replicate). **(b)** Number of cells per replicate from cell line HEL11.4. **(c)** Percentage of cells unmapped in the first annotation step or annotated as panRG/ExPanNeu (within unmapped cells) in the second annotation step, from cell line HEL11.4.

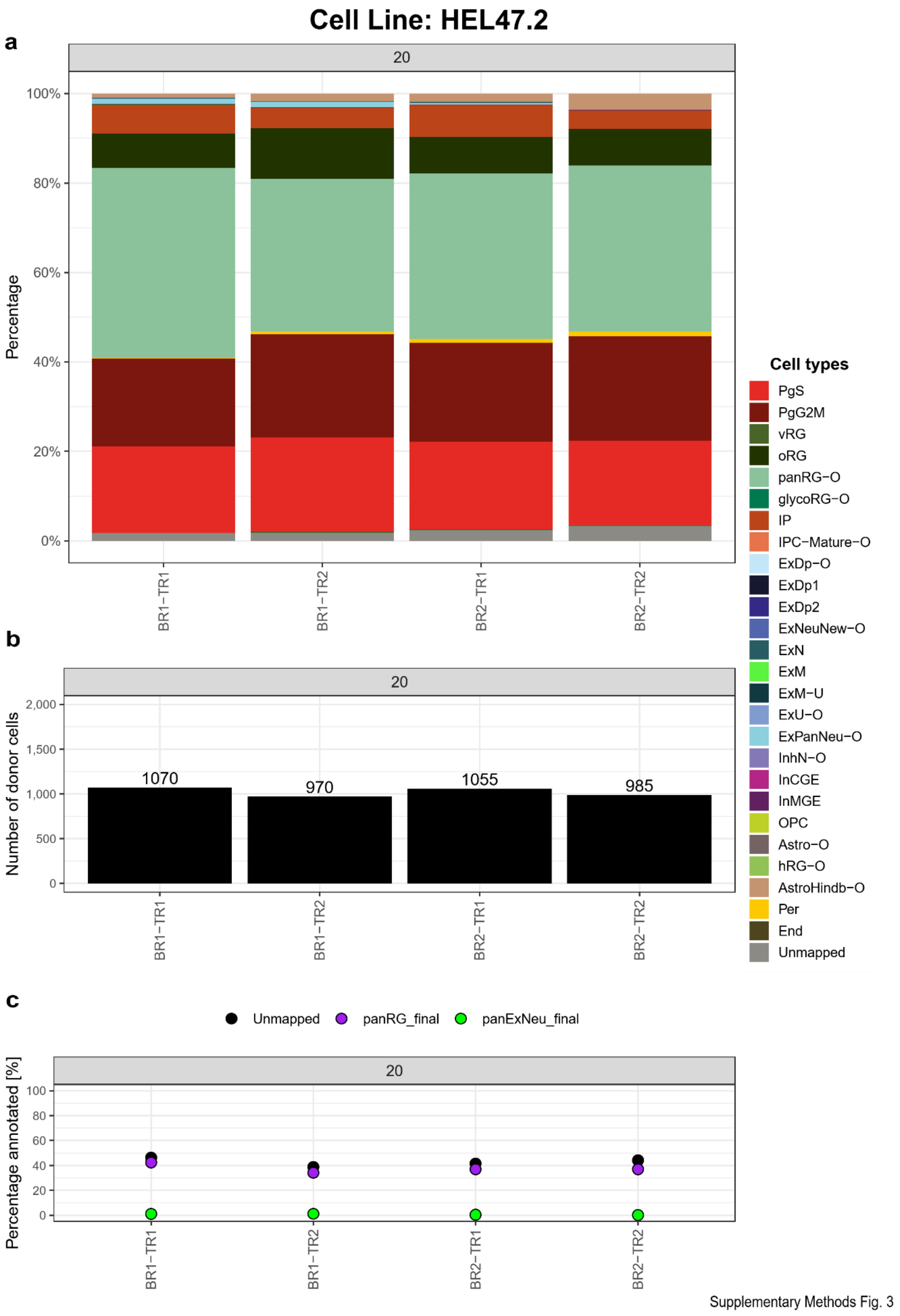


**Supplementary Methods Fig. 3. Reproducibility of replicates in scRNA-seq: HEL47.2.** Same as in Supplementary Methods Fig. 2, for cell line HEL47.2.

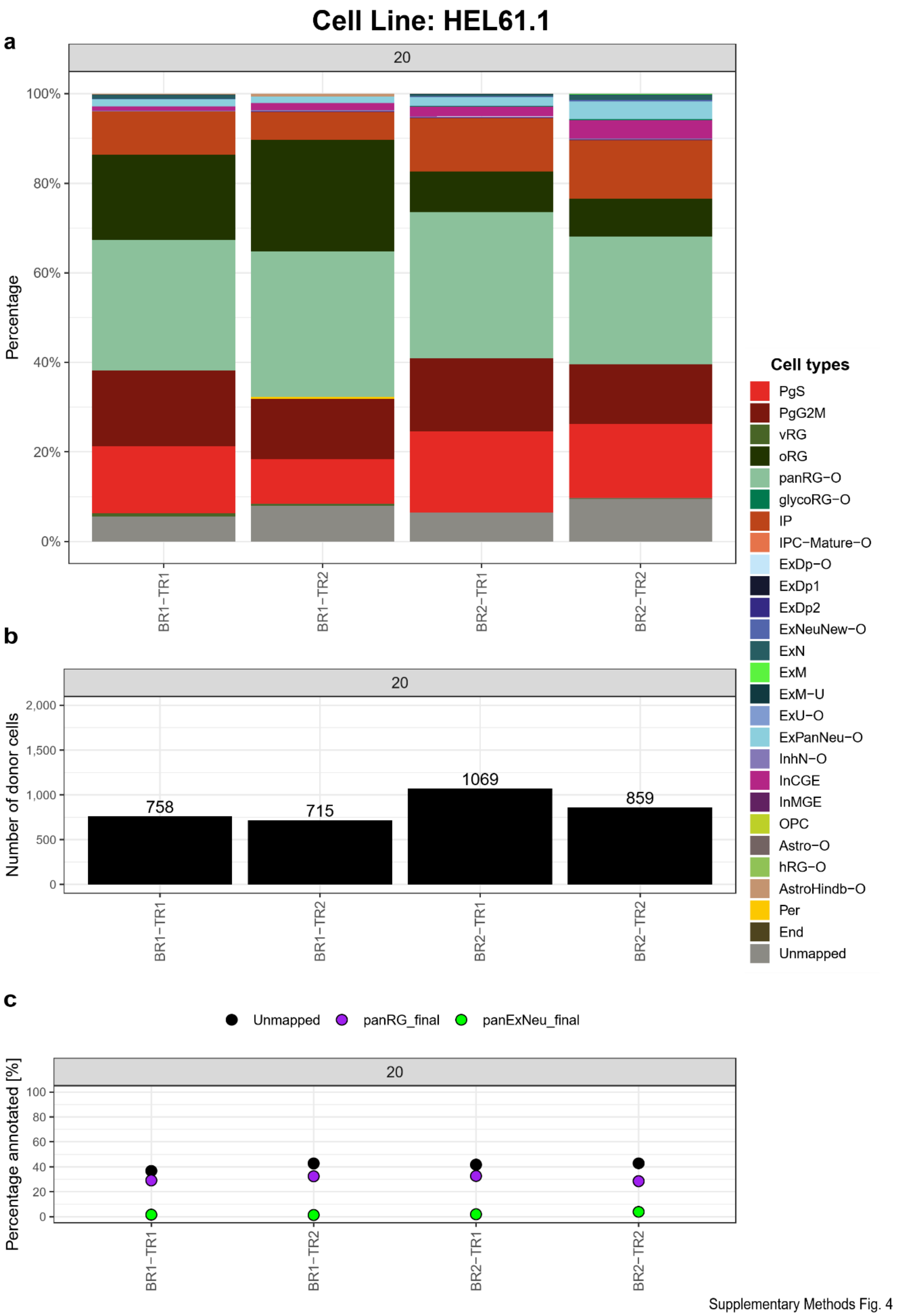


**Supplementary Methods Fig. 4. Reproducibility of replicates in scRNA-seq: HEL61.1.** Same as in Supplementary Methods Fig. 2, for cell line HEL61.1.

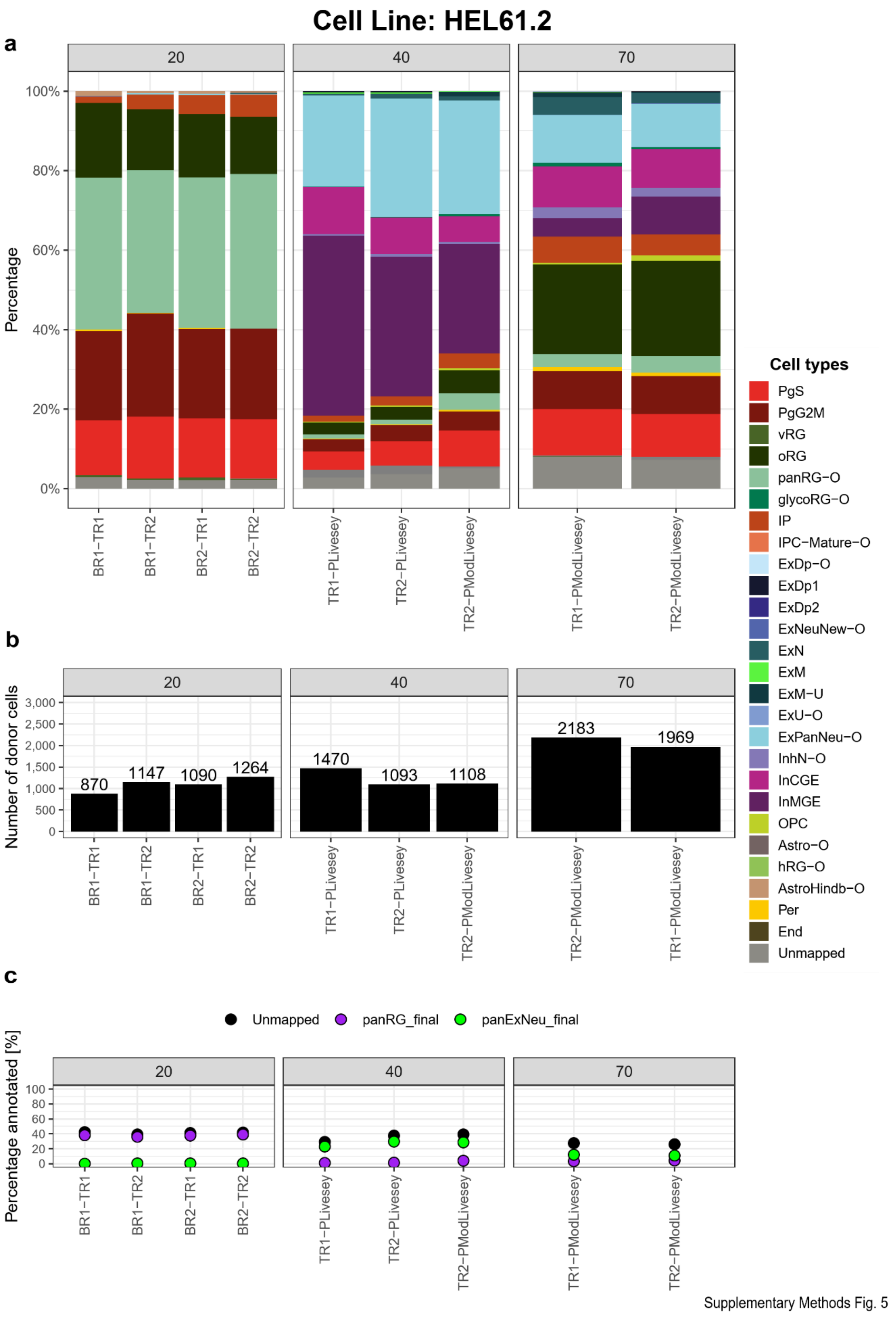


**Supplementary Methods Fig. 5. Reproducibility of replicates in scRNA-seq: HEL61.2.** Same as in Supplementary Methods Fig. 2, for cell line HEL61.2.

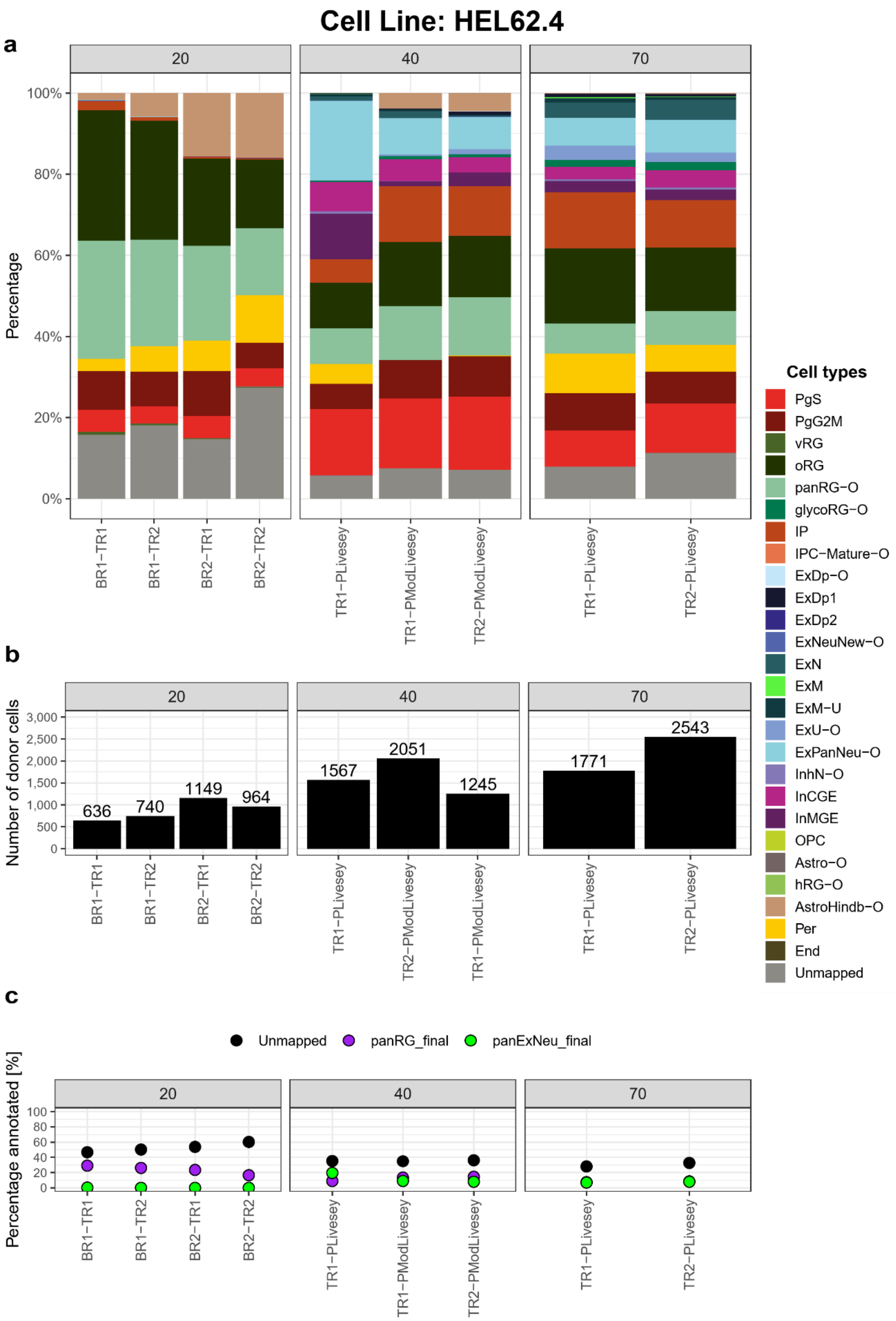


Supplementary Methods Fig. 6

**Supplementary Methods Fig. 6. Reproducibility of replicates in scRNA-seq: HEL62.4.** Same as in Supplementary Methods Fig. 2, for cell line HEL62.4.

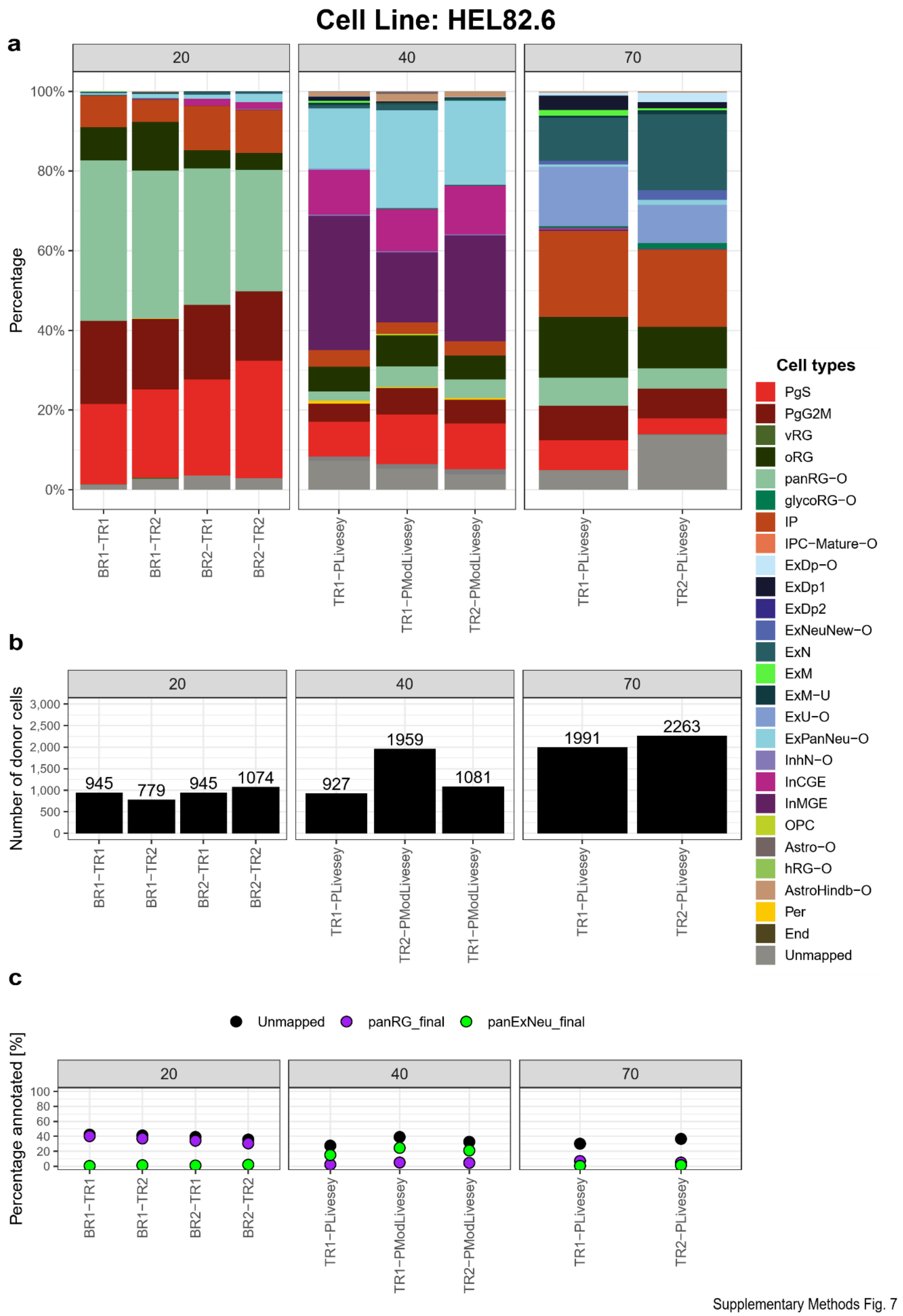


**Supplementary Methods Fig. 7. Reproducibility of replicates in scRNA-seq: HEL82.6.** Same as in Supplementary Methods Fig. 2, for cell line HEL82.6.

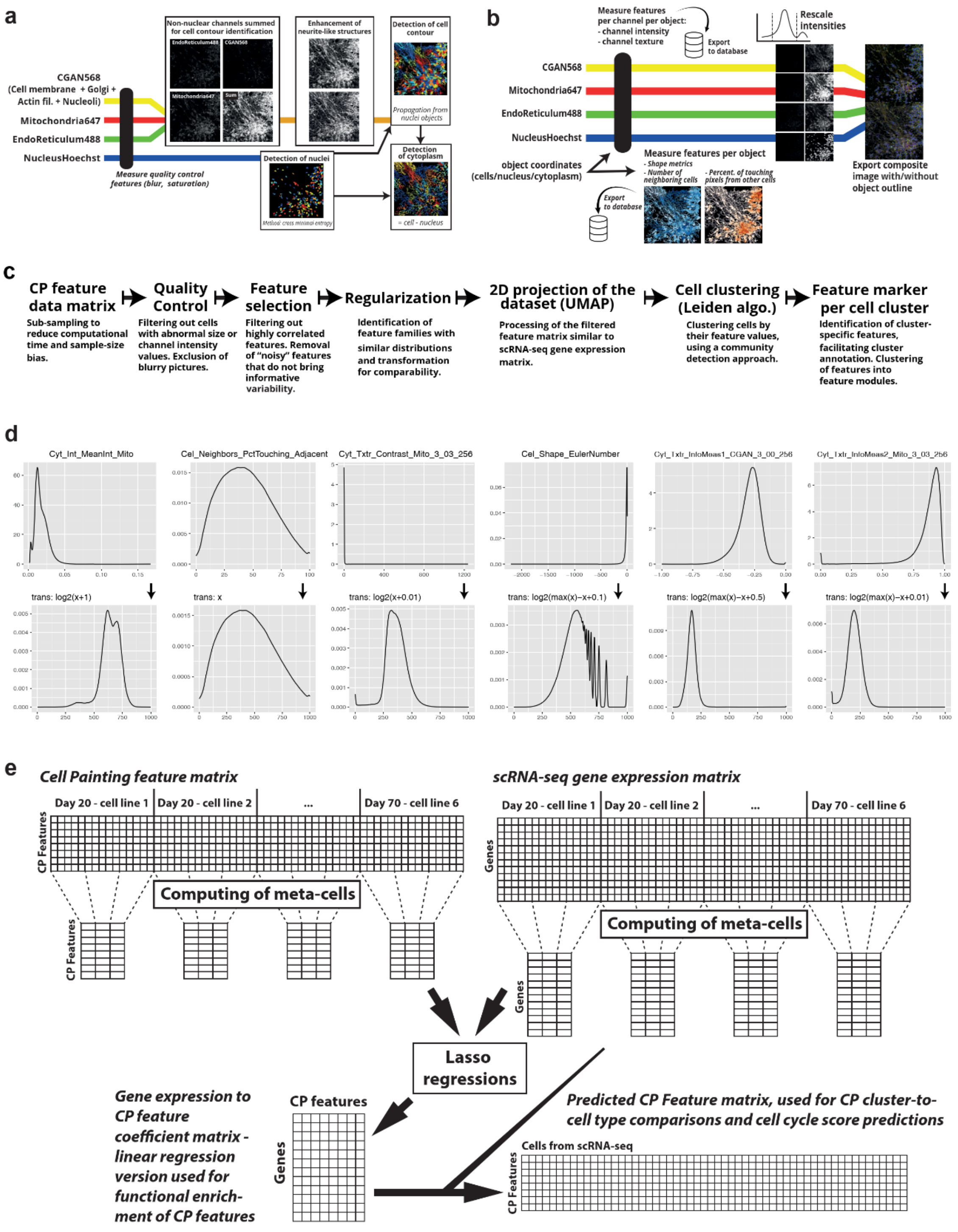


**Supplementary Methods Fig. 8. Cell Painting analysis workflow. (a)** Schematic of the CellProfiler segmentation steps. The nucleus channel was used to segment the nuclei, the three cytoplasmic channels were then summed to detect cell contours, by propagation from the nuclei. **(b)** Schematic of feature extraction steps and image export. **(c)** Secondary analysis workflow (processing of the Cell Painting feature matrix). **(d)** Effect of regularization on the CP feature distribution. Original distribution (first row) and regularized distribution (second row) are shown for one feature per family of transformations. **(e)** Schematic of the steps used to link gene expression to Cell Painting features. Metacells were computed from each experimental population and used to train lasso regression models, one per CP feature. The computed coefficients and intercepts were stored in a coefficient matrix, used for CP feature functional enrichment analysis. The coefficient matrix was also used with the gene expression data to predict the CP feature values of each cell from the scRNA-seq dataset.
