## Supplementary Fig for "Joint profiling of cell morphology and gene expression during *in vitro* neurodevelopment"

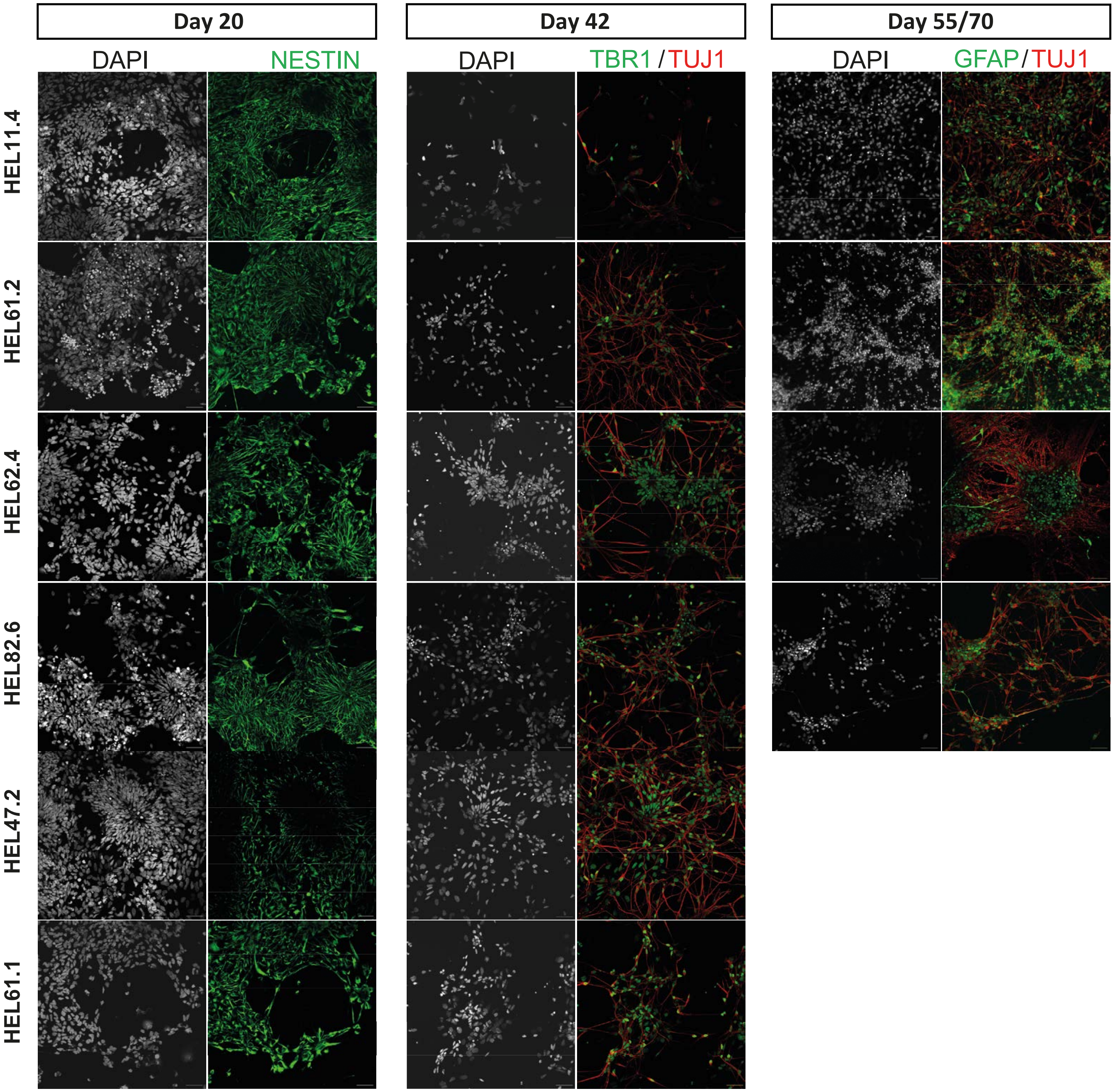


**Supplementary Figure 1: Related to figure 1, benchmarking of *in vitro* cell types.** Representative ICC images per donor showing canonical marker expression per time point.

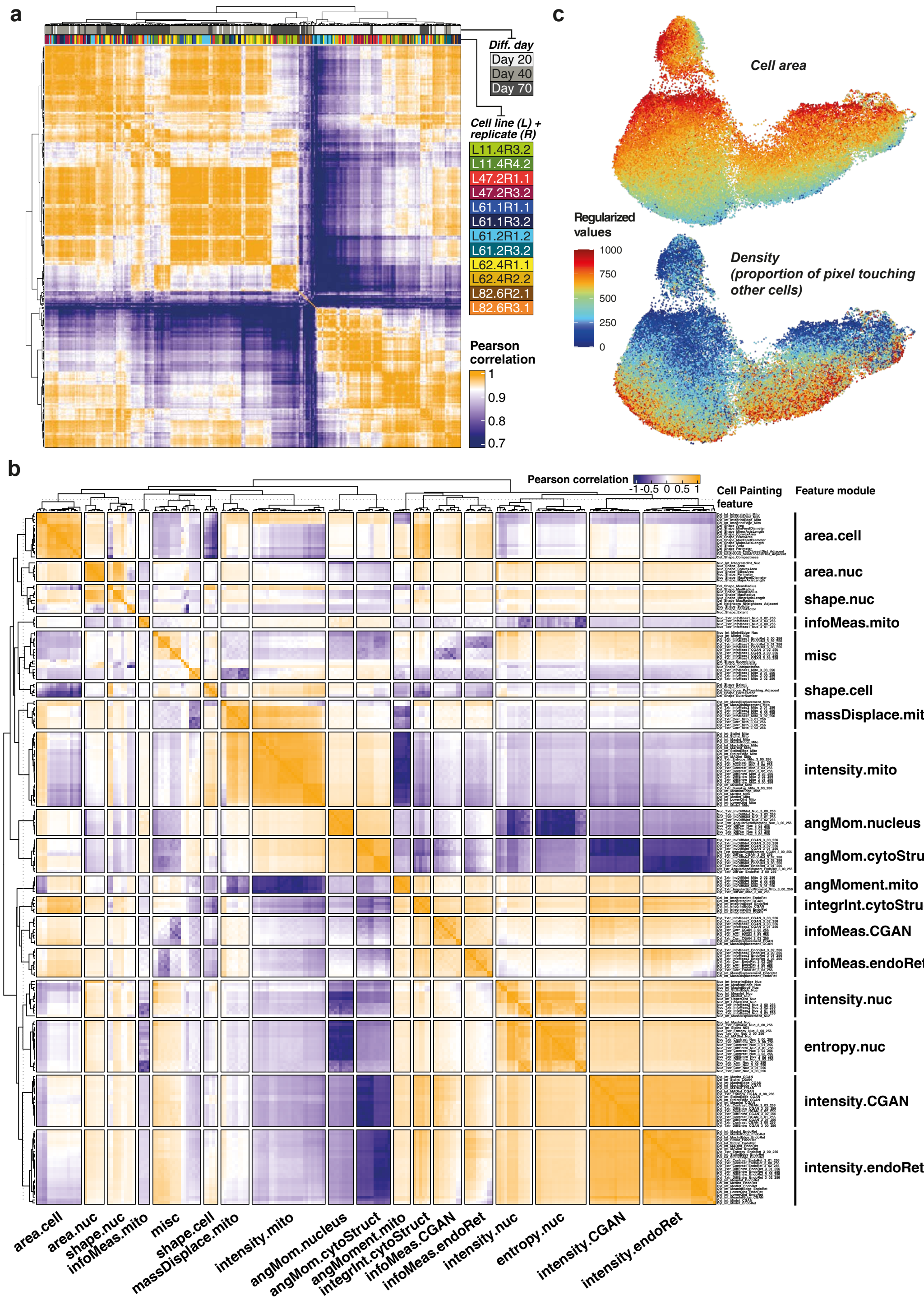


**Supplementary Figure 2: Related to figure 2, additional Cell Painting results. (a)** Well-to-well Pearson correlation of the Cell Painting (CP) dataset. Heatmap header annotates the corresponding differentiation day (first row) and the cell line-replicate combination (second row), per well. **(b)** CP feature-to-feature Pearson correlation, grouped by CP feature module. The 'misc' module corresponds to unclassified CP features. The feature name nomenclature is as follows: [Compartment]_[Feature family]_[Feature name]_[Channel]_[Texture variation], where 'Compartment' indicates where the feature was measured (Cell, Cytoplasm or Nucleus); 'Feature family' describes the broad measure type of the feature (intensity, texture, shape, neighbors); 'Feature name' abbreviates the precise name of the measure (for example, MedInt is median intensity); 'Channel' indicates which channel the feature was extracted from (only for intensity or texture features); and 'Texture variation' provides information about how the texture was extracted (only for texture features). **(c)** CP regularized features values corresponding to Cell area (Cel_Shape_Area) and density (Cel_Neighbors_PctTouching_Adjacent) projected on the CP UMAP.

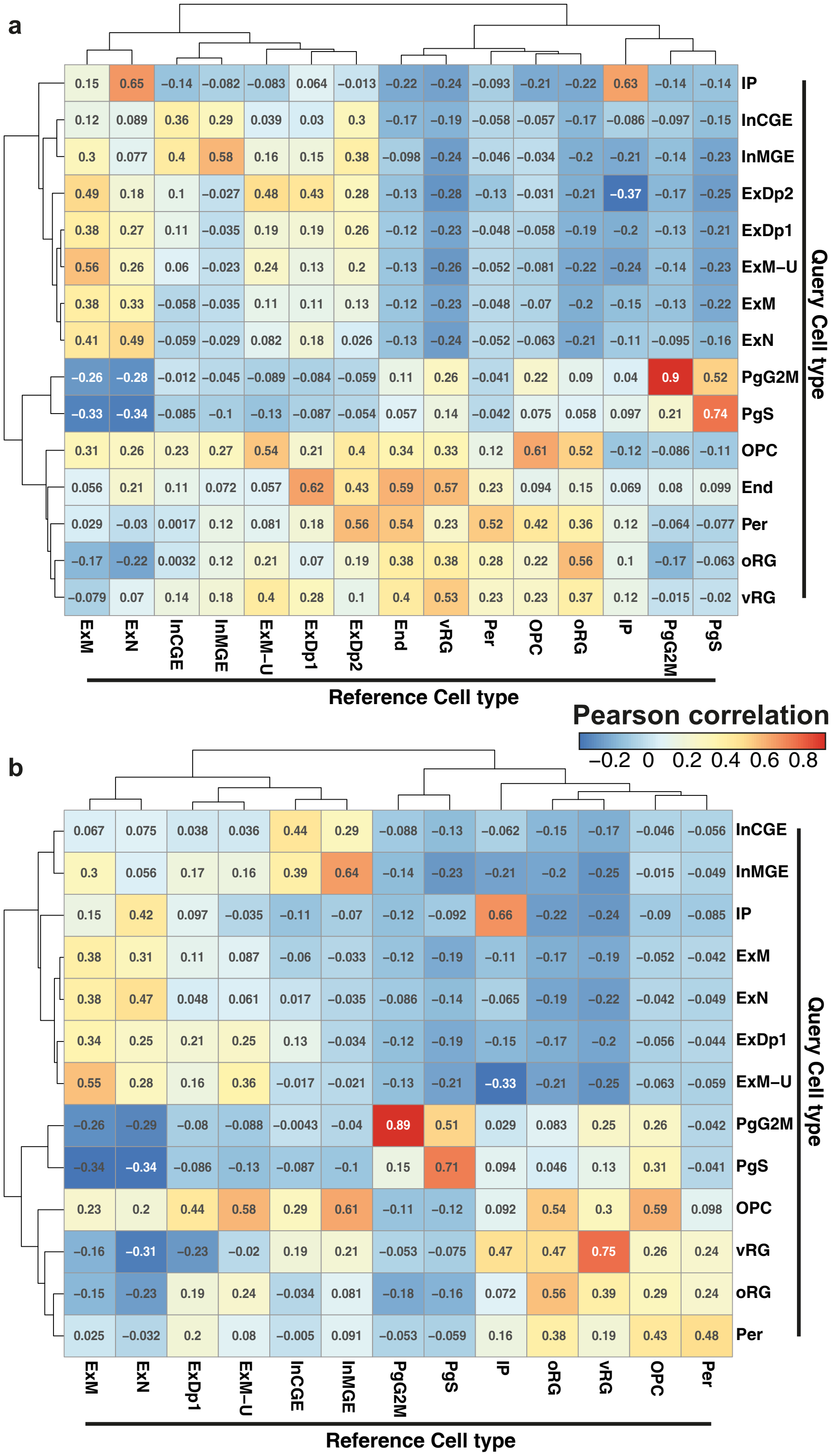


**Supplementary Figure 3: Related to figure 2, *in vitro* to fetal cell type correlations. (a)** Correlation of marker gene expression between annotated *in vitro* (query) cell types and fetal (reference) cell types, prior to filtering out unmapped cells (mean R=0.50, off-diagonal mean R=0.05). **(b)** Same correlation as in (a), after filtering out unmapped cells showing higher specificity (mean R=0.55, off-diagonal mean R=0.03).

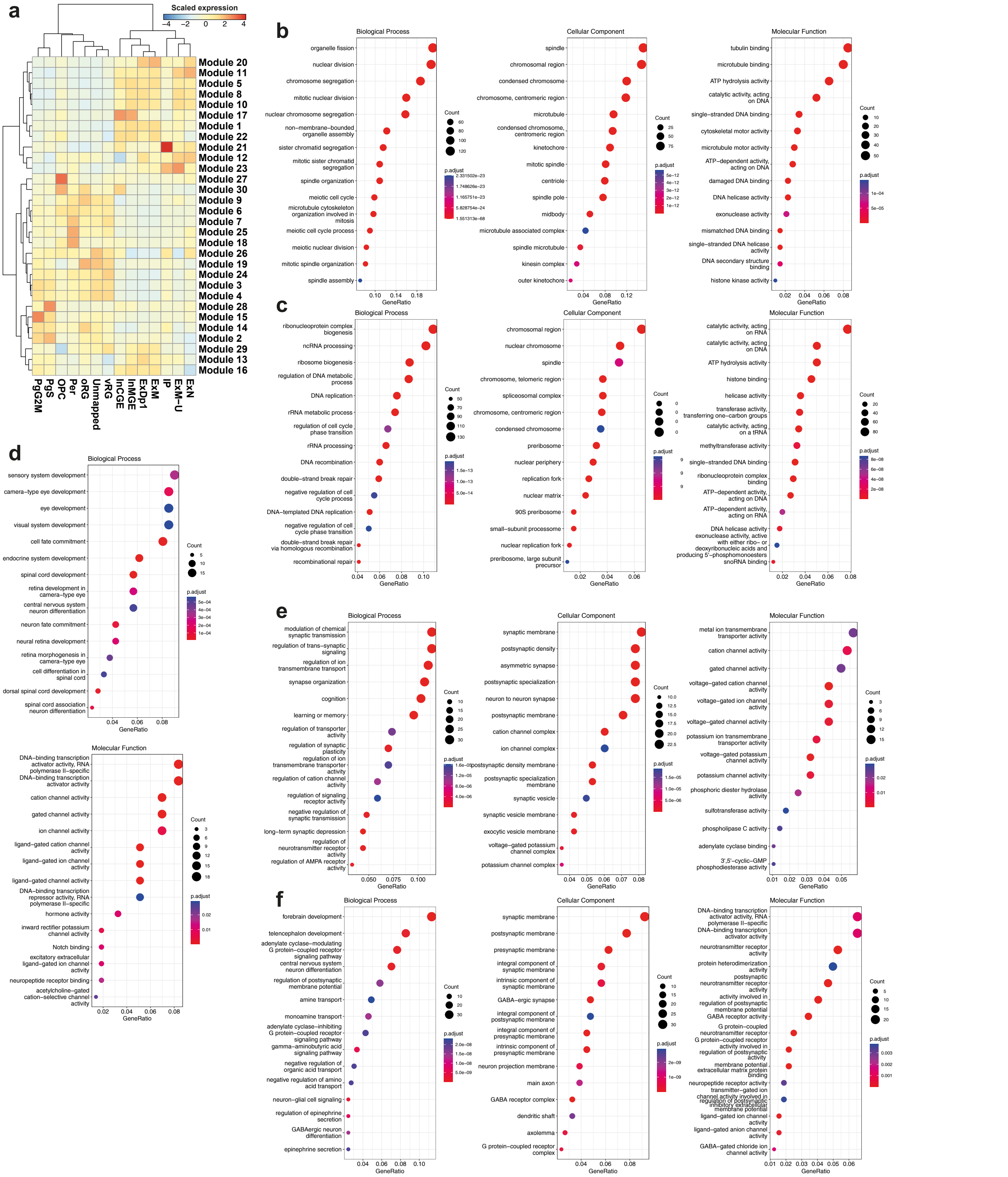

**Supplementary Figure 4: Related to figure 3, functional enrichment across the developmental trajectory.** **(a)** Aggregated expression per cell type of modules of genes changing expression as a function of pseudotime. **(b-f)** Enrichment of GO terms (for each of the three sub-ontologies: 'Biological Process', 'Cellular Component', and 'Molecular Function') within the top-activated module for **(b)** PgG2M (module 15), **(c)** PgS (module 2), **(d)** IP (module 21), **(e)** ExM & ExDp1 (module 20), and **(f)** InMGE & InCGE (module 17). No significant terms were found for 'Cellular Component' within module 21 **(d)**.

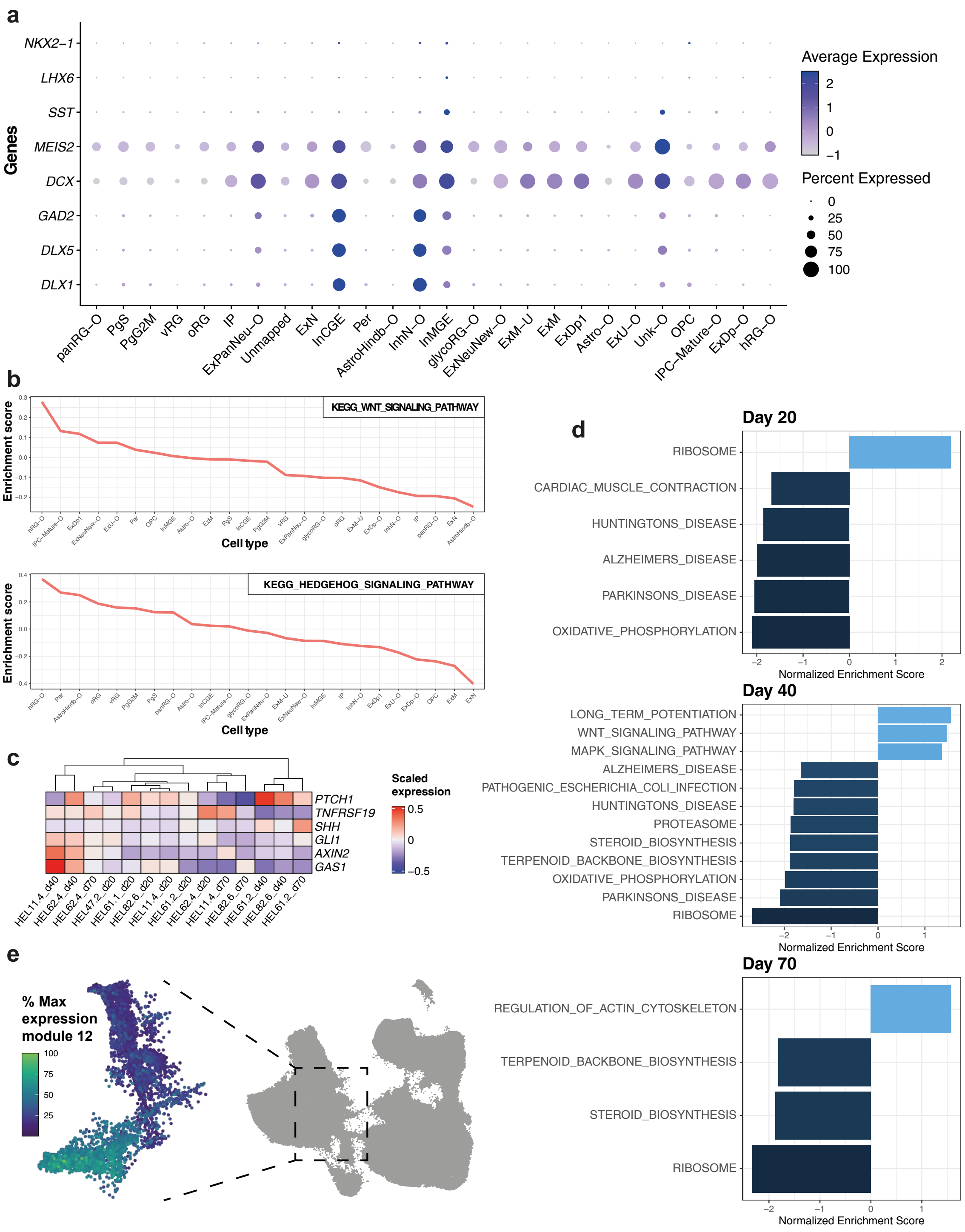


**Supplementary Figure 5: Related to figure 4, characterization of inhibitory neurons produced *in vitro*.(a)** Dot plot representing the expression of markers of CGE- and MGE-like interneurons across all cell types from the two-step annotation. **(b)** GSVA enrichment scores per pseudobulked cell type for WNT and Hedgehog signaling pathways, ordered from highest to lowest enrichment. **(c)** Scaled gene expression aggregated per experimental population (time-point × cell line) of genes representing key WNT/Hedgehog signaling pathway readouts as per Strano et al. (2020)[31]. 'Ventralized' experimental populations (producing more inhibitory neurons, Fig. 5c) show higher *PTCH1* expression as previously reported[37], but other genes do not follow reported patterns (higher *PTCH1* and *GLI1*, and lower *GAS1* in ventralized differentiations; higher *AXIN2* and *TNFRSF19* in dorsalized differentiations). **(d)** GSEA (KEGG) of differentially expressed genes between inhibitory neurons (InMGE, InCGE & InhN-O) and excitatory neurons (ExDp-O, ExM-U, ExN, ExNeuNew-O, ExDp1 ,ExM, ExU-O), at each differentiation time point. **(e)** Representation of the excitatory-inhibitory branching segment within the pseudotime trajectory, as compared to the overall UMAP (shown in gray), and the scaled expression of the inhibitory branch-associated module (module 12) within this segment.

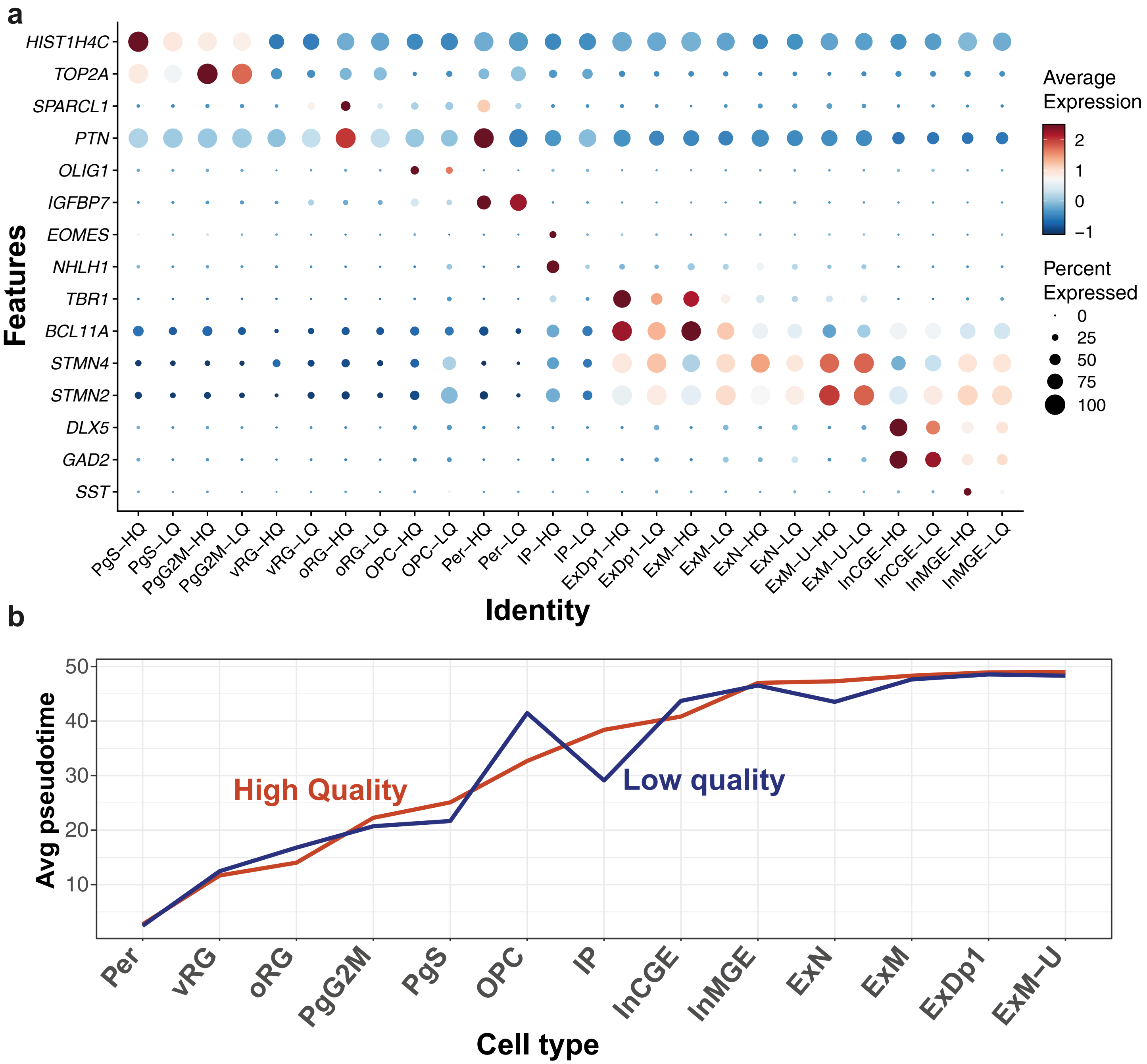


**Supplementary Figure 6: Related to figure 5, LQ and HQ cell type subsets show similarities in expression and pseudotime. (a)** Dot plot representing the expression of cell-type specific canonical markers in the HQ and LQ subsets of each cell type. **(b)** Average pseudotime values per cell type grouped by HQ and LQ subsets, ordered by increasing pseudotime of the HQ subset.

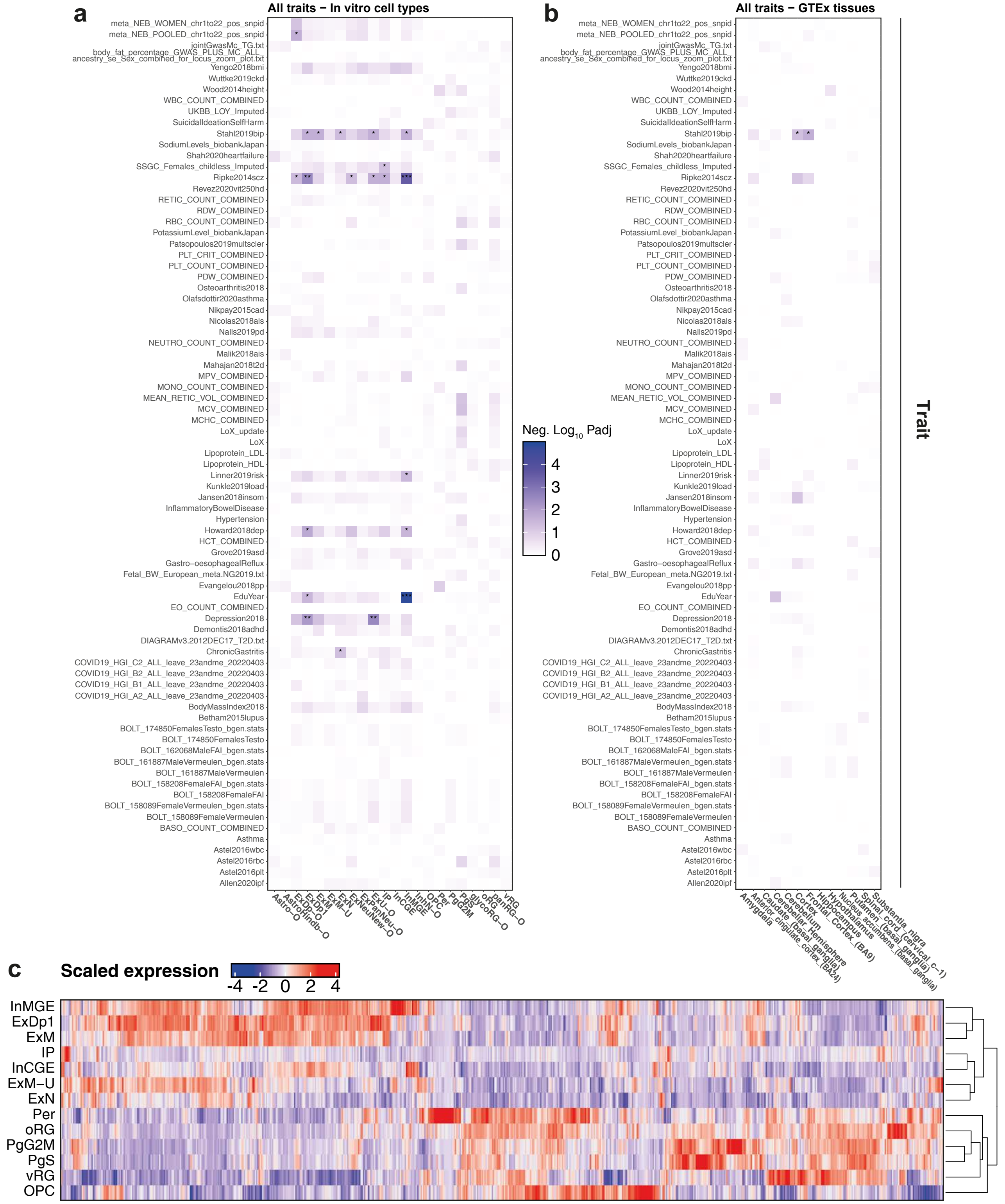


**Supplementary Figure 7: Related to figure 6, cell type-specificity of brain- and non-brain-related traits.** **(a)** Stratified LD score regression analysis shown for all analyzed traits (n=79) per cell type from our *in vitro* differentiation. Tile color represents the corresponding p values after multiple testing correction across traits, within cell types, with the significance level indicated as follows: pAdj<0.05(*), pAdj<0.01(**) and pAdj<0.001 (***). **(b)** Stratified LD score regression analysis as in (a) per GTEx Brain tissue type. **(c)** Heatmap of normalized expression, pseudobulked per cell type and scaled per gene, across brain-specific genes associated with developmental disorders (DDD study). All DDD genes with the confidence category 'definitive' are shown (n=719). ExDp1, ExM, ExM-U, ExN, InCGE and InMGE together were considered as mature neuronal cell types, whereas PgS, PgG2M, oRG and vRG were considered progenitors.
