## Supplementary figures and images for "Joint profiling of cell morphology and gene expression during *in vitro* neurodevelopment"

### Fig. S1

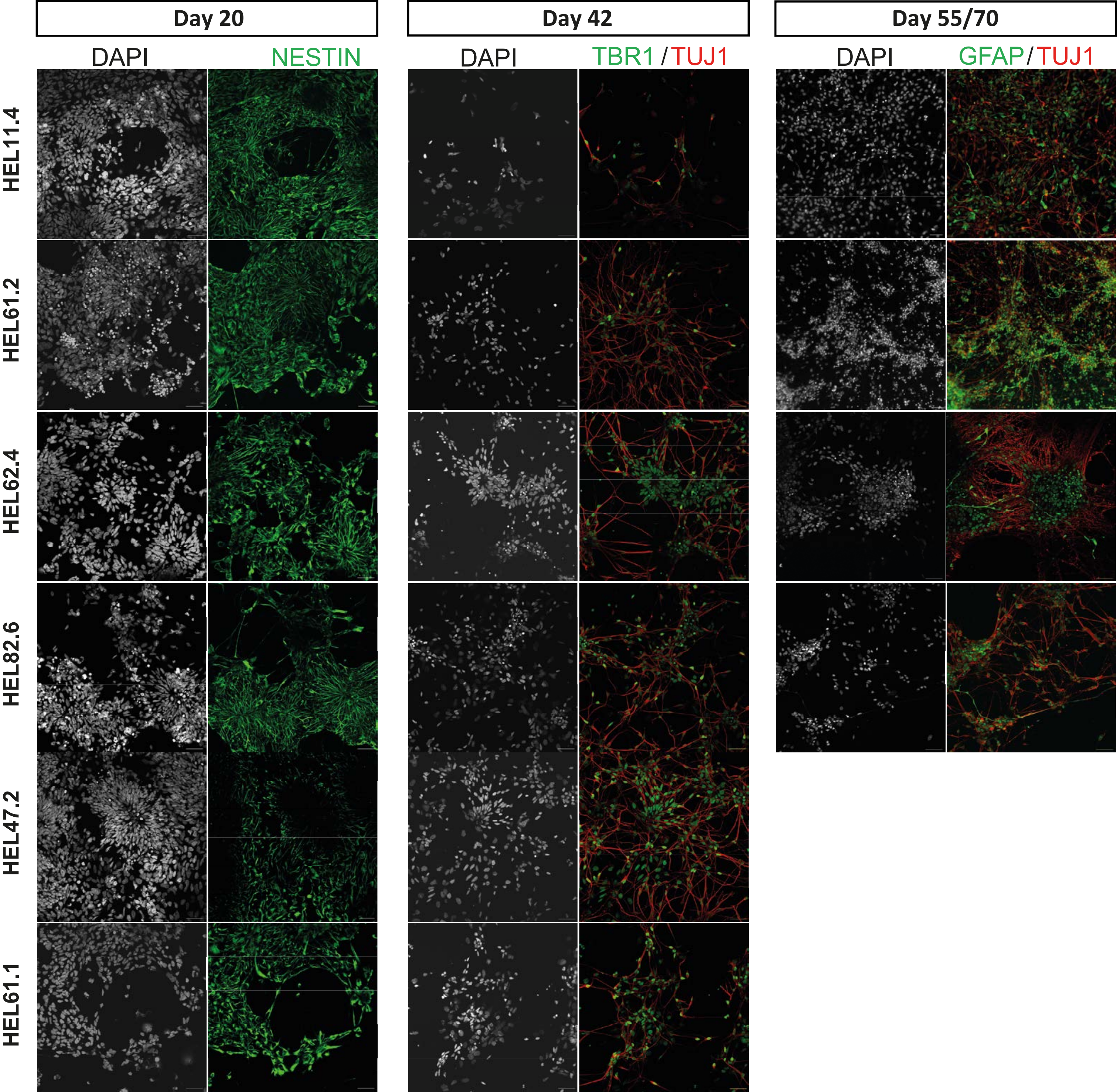

### Fig. S2

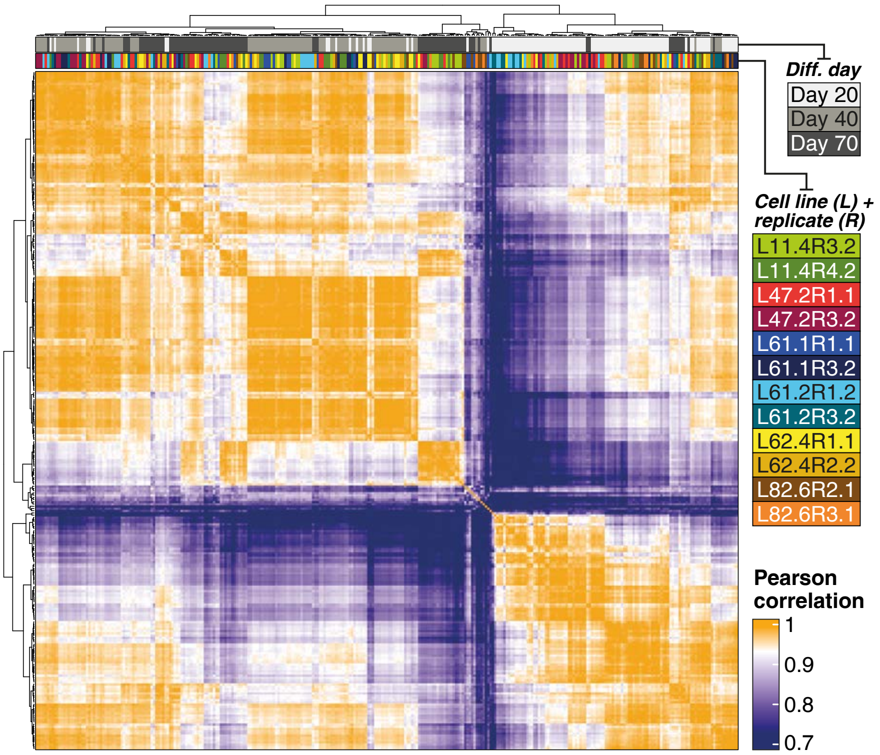

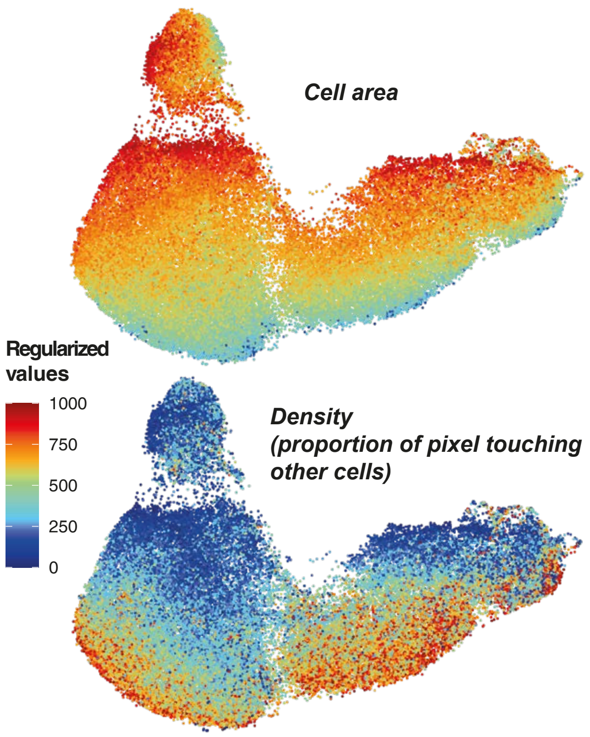

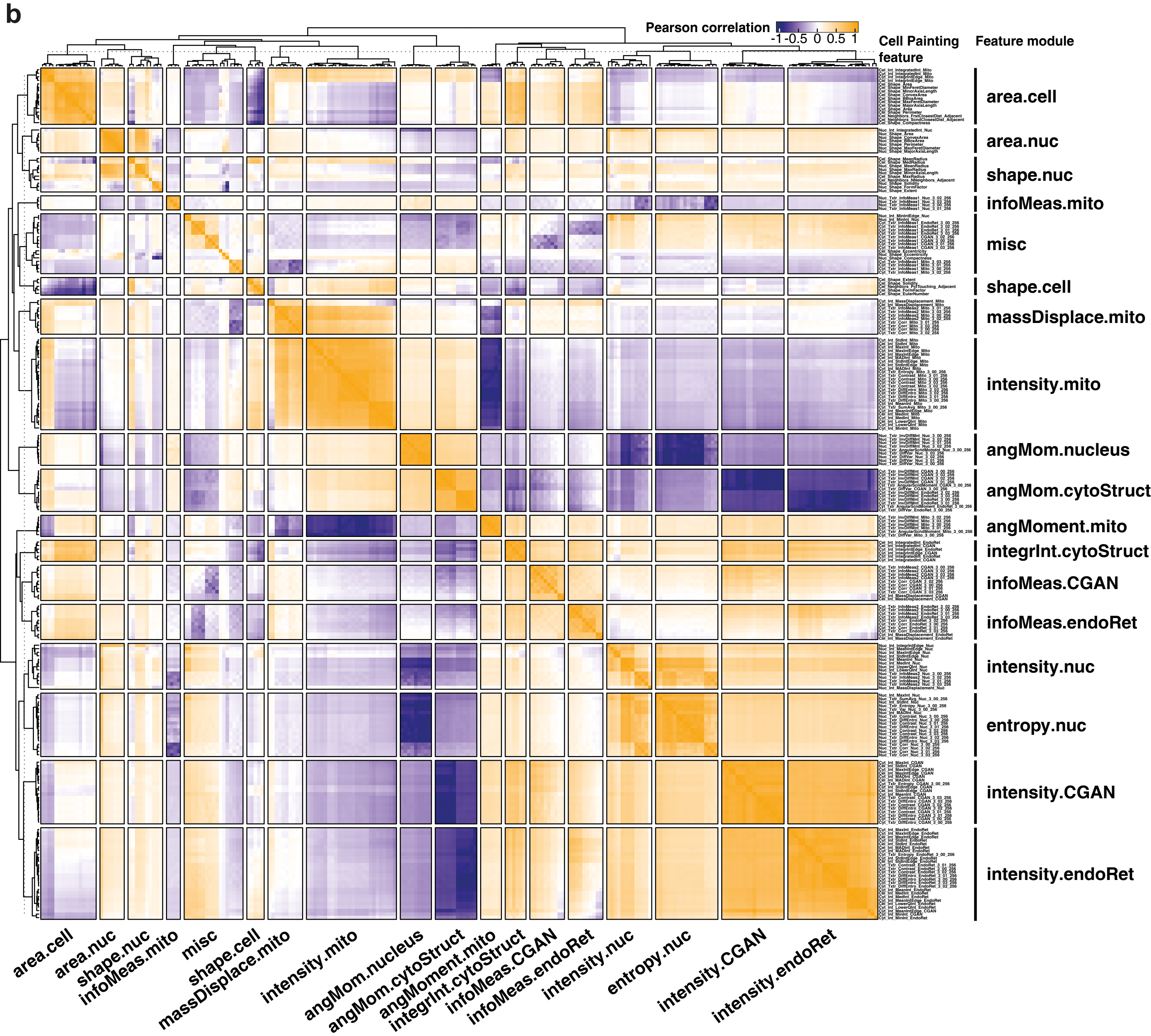

### Fig. S3

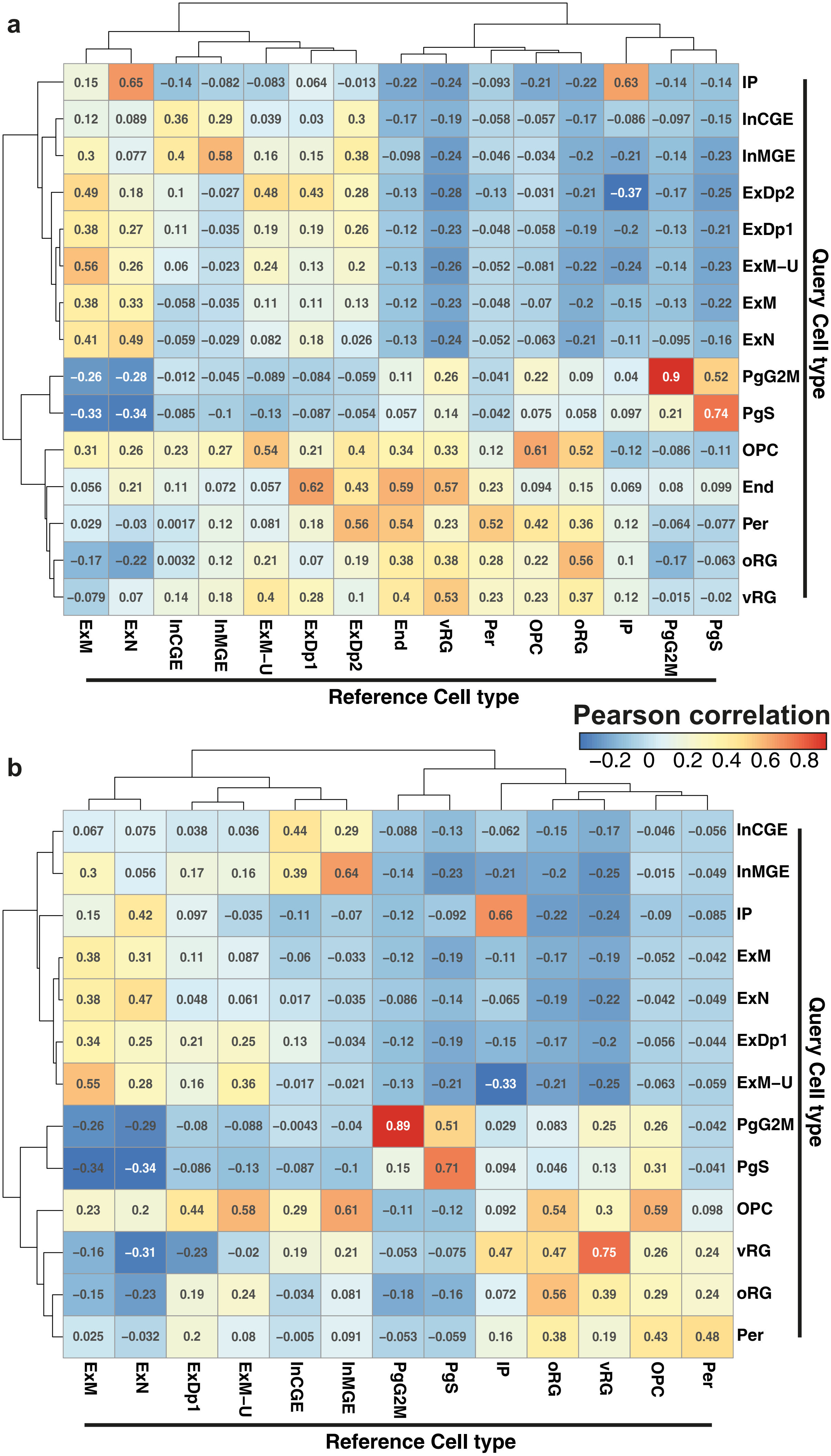

### Fig. S4

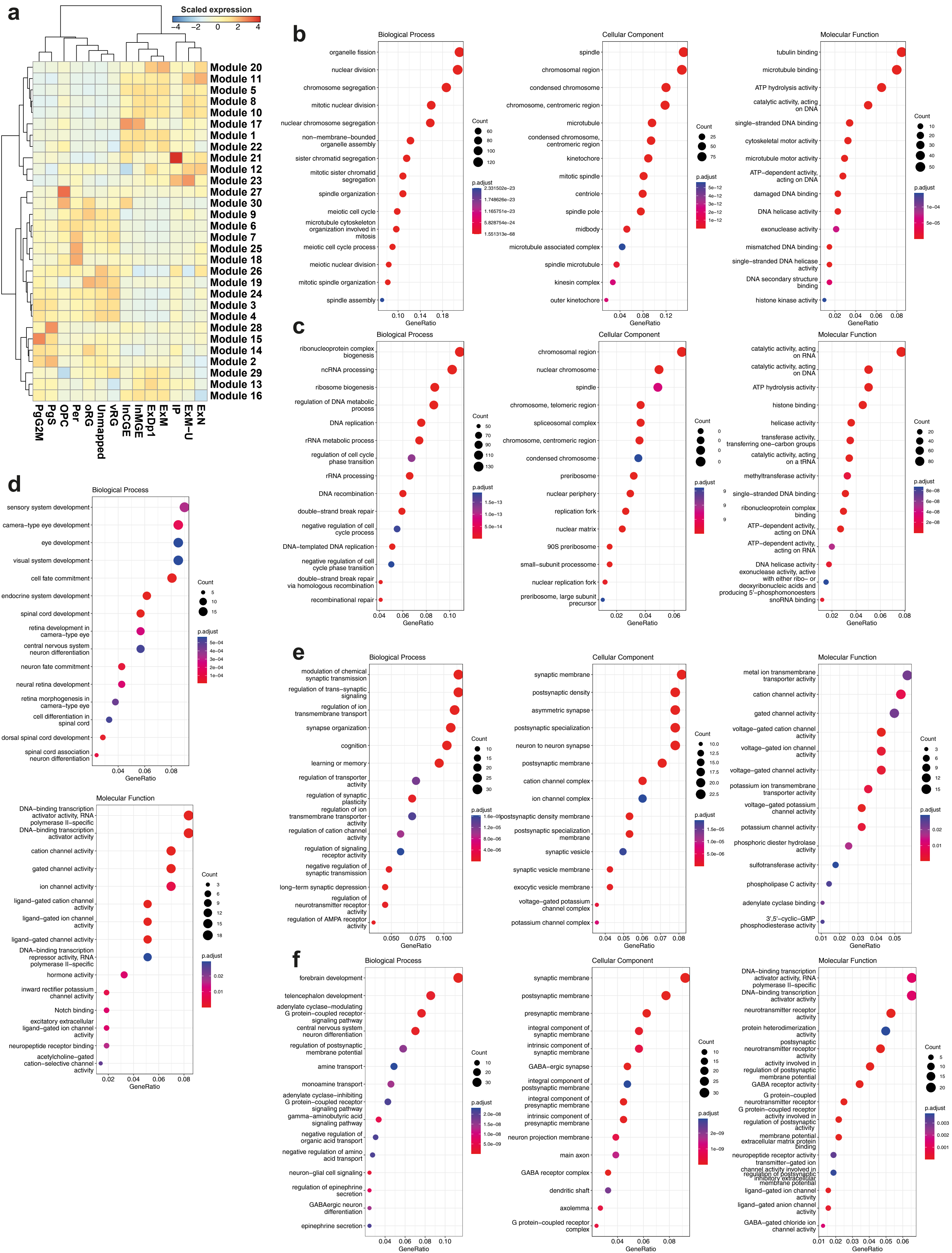

### Fig. S5

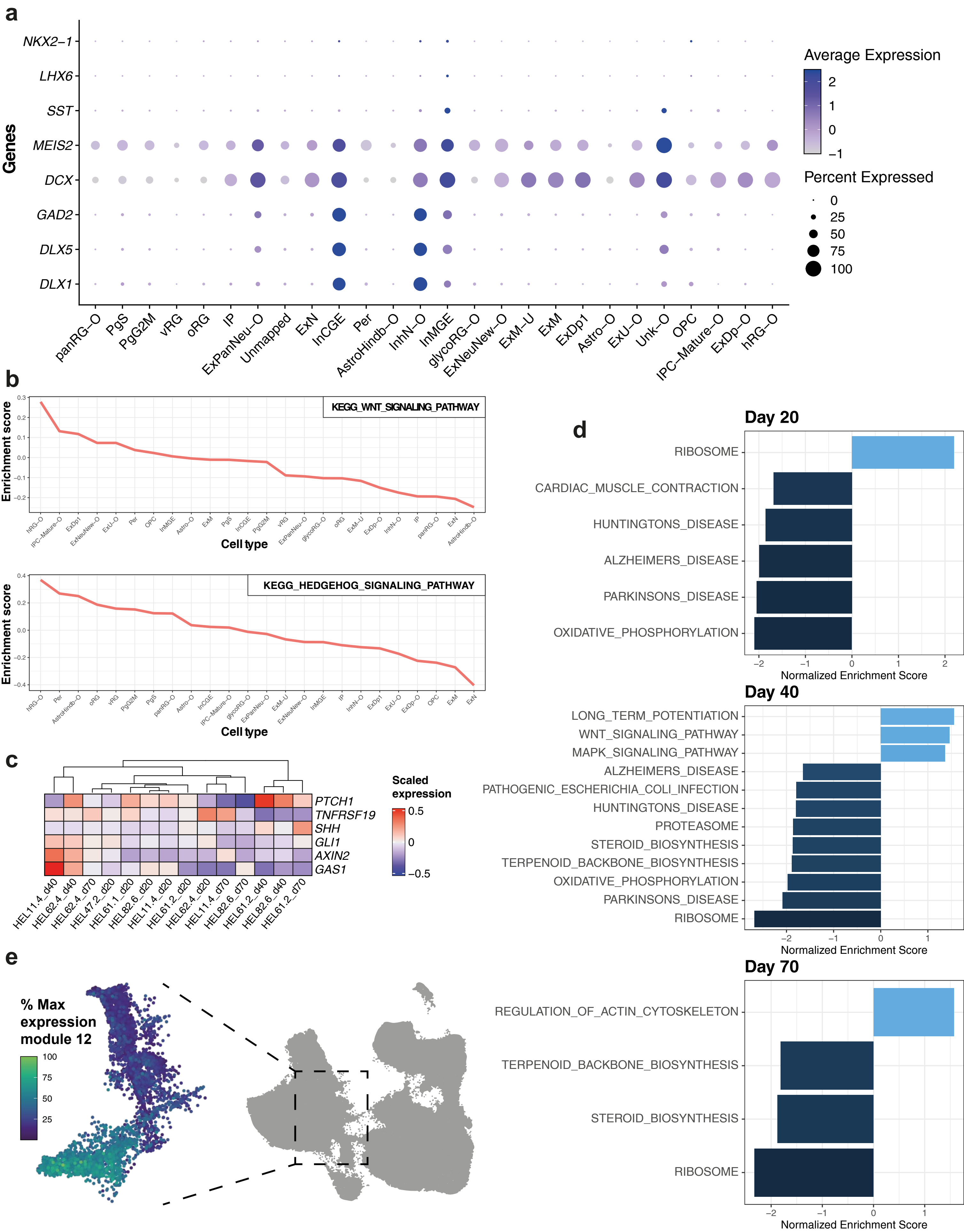

### Fig. S6

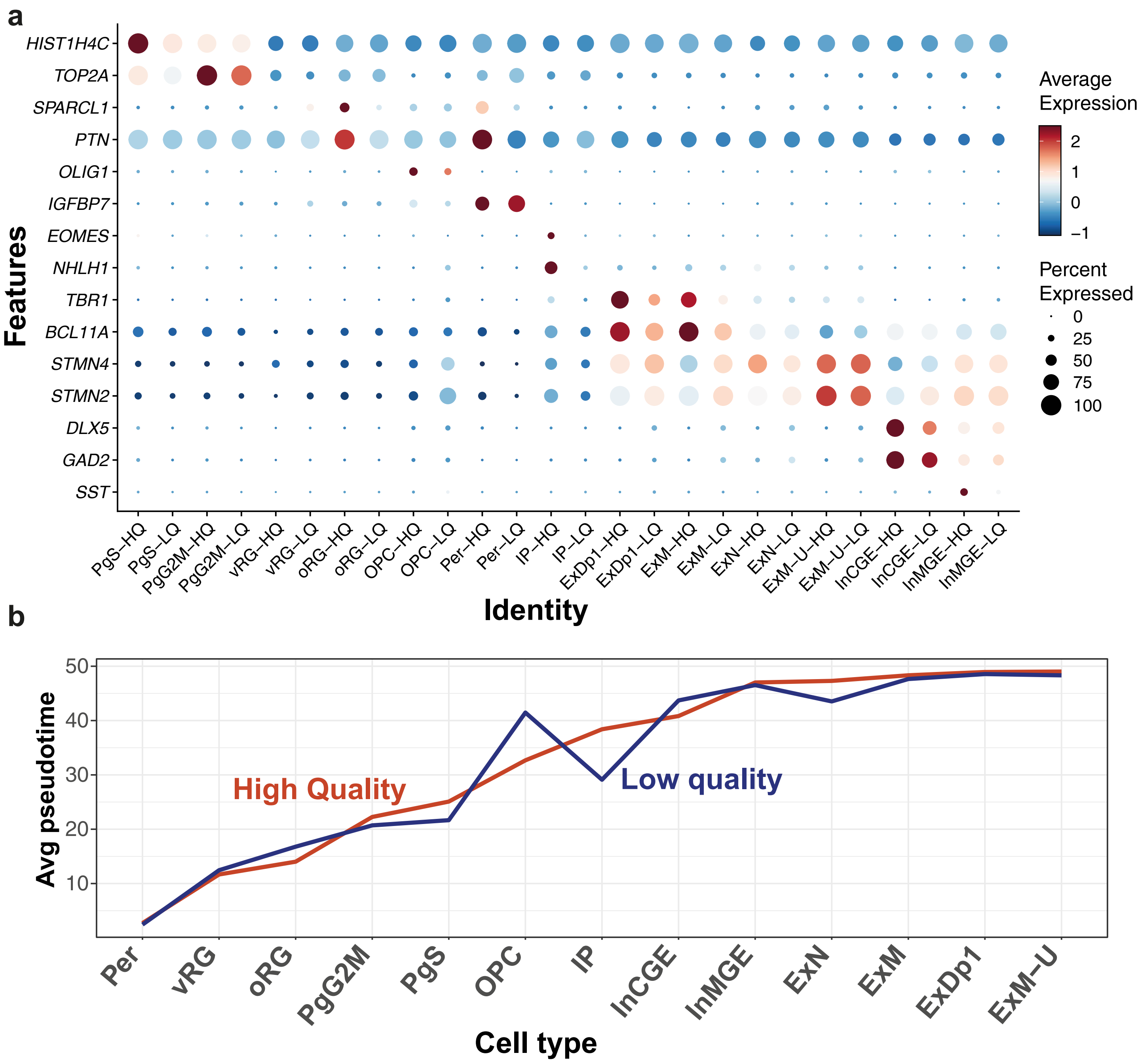

### Fig. S7

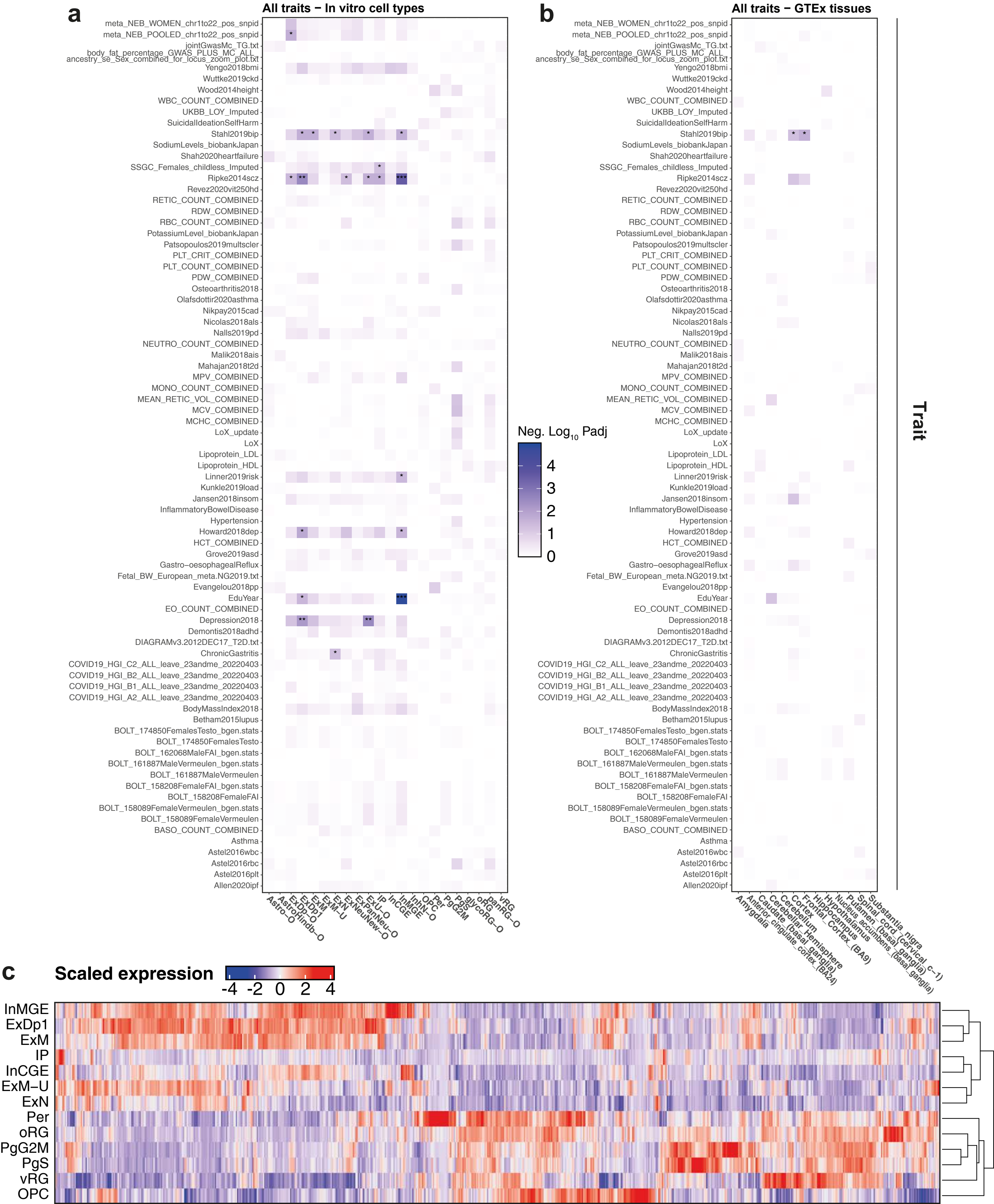
